# Mouse embryonic stem cells self-organize into trunk-like structures with neural tube and somites

**DOI:** 10.1101/2020.03.04.974949

**Authors:** Jesse V Veenvliet, Adriano Bolondi, Helene Kretzmer, Leah Haut, Manuela Scholze-Wittler, Dennis Schifferl, Frederic Koch, Milena Pustet, Simon Heimann, Rene Buschow, Lars Wittler, Bernd Timmermann, Alexander Meissner, Bernhard G Herrmann

**Affiliations:** Dept. Of Developmental Genetics, Max Planck Institute for Molecular Genetics, Ihnestr. 63-73, 14195 Berlin, Germany; Dept. Of Genome Regulation, Max Planck Institute for Molecular Genetics, Ihnestr. 63-73, 14195 Berlin, Germany; Microscopy and Cryo-Electron Microscopy, Max Planck Institute for Molecular Genetics, Ihnestr. 63-73, 14195 Berlin, Germany; Sequencing Core Facility, Max Planck Institute for Molecular Genetics, Ihnestr. 63-73, 14195 Berlin, Germany; Institute for Medical Genetics, Charité - University Medicine Berlin, Campus Benjamin Franklin, Hindenburgdamm 30, 12203 Berlin, Germany

## Abstract

Post-implantation embryogenesis is a highly dynamic process comprising multiple lineage decisions and morphogenetic changes inaccessible to deep analysis *in vivo*. Mouse embryonic stem cells (mESCs) can form aggregates reflecting the post-occipital embryo (*gastruloids*), but lacking proper morphogenesis. Here we show that embedding of aggregates derived from mESCs in an extracellular matrix compound results in Trunk-Like-Structures (TLS) with a high level of organization comprising a neural tube and somites. Comparative single-cell RNA-seq analysis demonstrates that TLS execute gene-regulatory programs in an embryo-like order, and generate primordial germ cell like cells (PGCLCs). TLS lacking *Tbx6* form ectopic neural tubes, mirroring the embryonic mutant phenotype. ESC-derived trunk-like structures thus constitute a novel powerful *in vitro* platform for investigating lineage decisions and morphogenetic processes shaping the post-implantation embryo.

**One sentence summary:** A platform for generating trunk-like-structures with precursors of spinal cord, bone and muscle from stem cells in a dish

## Background

Vertebrate post-implantation development comprises a multitude of complex morphogenetic processes, which result from self-organization of stem cells and their descendants shaping the embryonic body plan^1^. Recently developed stem cell models represent powerful platforms for deconstructing the dynamics of these processes that are inaccessible in the embryo^1, 2^. The most advanced models in terms of developmental stage accomplished so far are gastruloids^2, 3^, aggregates of mESCs able to self-organize into the three body axes^3^. However, gastruloids lack proper morphogenesis, such as a failure of neural tube formation from neural cells or somite condensation from presomitic cells^3^. *In vivo*, the extracellular matrix (ECM) has a critical role in tissue morphogenesis^4^. *In vitro*, matrigel can serve as ECM surrogate, and culture media supplemented with a low percentage of matrigel have been shown to induce complex morphogenesis in organoids^5^.

We therefore explored if embryo-like morphogenetic features could be induced by embedding 96h post aggregation gastruloids in 5% matrigel for an additional 24h period (**Fig. 1a**). We also tested a combination of matrigel together with the WNT signaling activator CHIR99021 (CHIR) and the BMP signaling inhibitor LDN193189 (LDN), that have been reported to induce a (pre-)somitic mesoderm ((P)SM) fate in 2D and 3D differentiation protocols^6–8^. To facilitate high-throughput characterization and quantification of our conditions, we generated mESCs with *T::H2B-mCherry* (hereafter T^mCH^) and *Sox2::H2B-Venus* (hereafter Sox2^VE^) reporters, marking the mesodermal (ME) or neural (NE) lineage respectively (**Extended Data Fig. 1a**). Strikingly, embedding in 5% matrigel alone was sufficient for segmentation in the T^mCH+^ ME domain and formation of a Sox2^VE+^ neural tube like structure (**Fig. 1b,d; Extended Data Fig. 1b,c**). The vast majority of structures (hereafter referred to as TLS for **T**runk-**L**ike-**S**tructures) elongated and formed a T^mCH+^ pole at the posterior end, with segmentation occurring in about half the TLS (**Fig. 1c**). Segments were T^mCH+^ demonstrating their mesodermal origin, and whole-mount *in situ* hybridization for Tcf15 and Uncx confirmed their somite identity (**Fig. 1d,e**)^7^. Additional CHIR treatment, both alone (hereafter TLS^C^) or in combination with LDN (hereafter TLS^CL^) improved the physical separation of neighboring segments without affecting T^mCH+^ pole formation or elongation (**Fig. 1b,c; Extended Data Fig. 1d-g**), and resulted in an excess of segments at the anterior end, arranged like a “bunch of grapes” (**Fig. 1b,d**)^9^. Moreover, the ME domain expanded at the expense of the NE compartment, with apparent disorganization of the posterior end and neural tissue (**Fig. 1b; Extended Data Fig. 1f,g**), as well as reduction of T^mCH+^/Sox2^VE+^ putative neuromesodermal progenitors (NMPs) - bipotent cells that give rise to both post-occipital NE and ME -, as confirmed by flow cytometry (**Fig. 1f**)^10^. In all three TLS protocols, segments were similar in size to embryonic somites (**Extended Data Fig. 1h**).

**Fig. 1.**
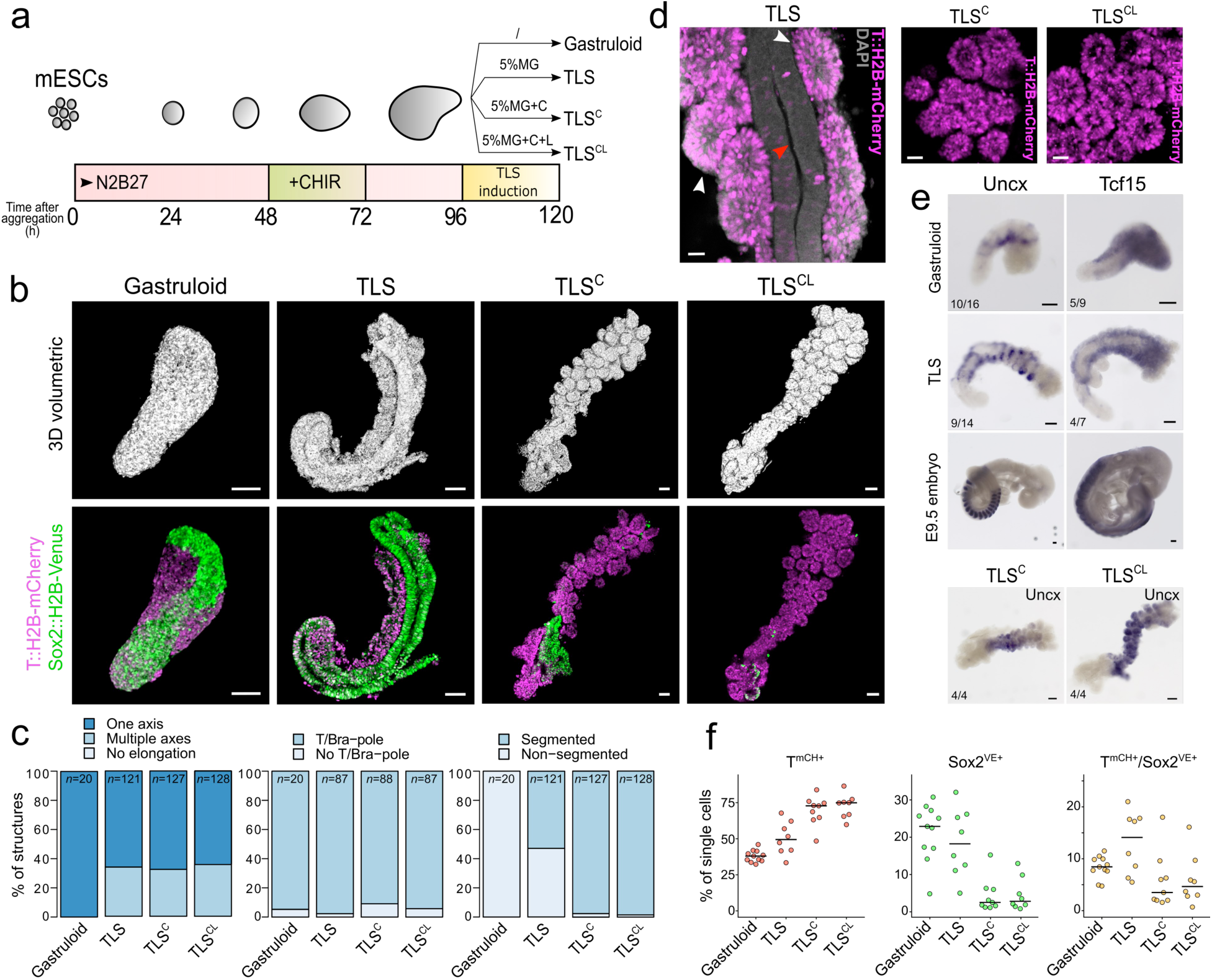
Generation of Trunk-Like Structures with somites and a neural tube. **a,** Schematic of the culture protocol: 200-250 mESCs were aggregated in ultra-low-attachment plates; *Wnt* agonist CHIR99021 (CHIR) was added between 48 and 72 hours (*3*). 96 aggregates were embedded in 5% matrigel (TLS). Optionally, structures were treated with CHIR alone (TLS^C^) or in combination with the BMP antagonist LDN (TLS^CL^). **b**, 3D volumetric renderings (upper panel) and confocal sections (bottom panel) of gastruloids, TLS, TLS^C^ and TLS^CL^. Scale bars 100μm. Each image is representative of at least ten biological replicates with similar morphology and expression patterns. **c**, Quantification of morphometric characteristics in gastruloids and TLS (see Supplemental Information for scoring criteria). **d,** Segments in TLS are T^mCH+^ and positioned adjacent to the neural tube in TLS. In TLS^C^ and TLS^CL^ the segments are arranged in “bunches of grapes”. Scale bars 25μm. Red arrowhead indicates neural tube, white arrowhead somites. **e**, Segments express somitic markers Tcf15 and Uncx as shown by whole-mount in situ hybridization. Note the characteristic stripy expression pattern of Uncx in TLS due to Uncx restriction to the posterior somite half, whereas Tcf15 is expressed throughout the segments (as in the embryo). Noteworthy, in TLS^C^ and TLS^CL^ Uncx is detected throughout the segments, indicating loss of anterior-posterior polarity. Scale bars 100μm. **f**, Percentage of T^mCH+^ (mesodermal), Sox2^VE+^ (neural), and T^mCH+/^Sox2^VE+^ (neuromesodermal) cells in gastruloids and TLS as measured by flow cytometry. Dots represent individual TLS/gastruloids, line indicates median.

In gastruloids, endodermal cells generally organize into a tubular structure resembling an embryonic gut^3^. Whole-mount immunofluorescence analysis of characteristic, endodermal expressed transcription factors FOXA2 and SOX17 also confirmed gut formation in our trunk-like structures (**Extended Data Fig. 2a-d**). Cells at the posterior base were SOX17-negative, but co-expressed FOXA2 with high levels of T^mCH^ (**Extended Data Fig. 2b**). Thus, our data show that embedding in matrigel is both necessary and sufficient to drive complex, embryo-like tissue morphogenesis of the three embryonic germ layers.

**Fig. 2.**
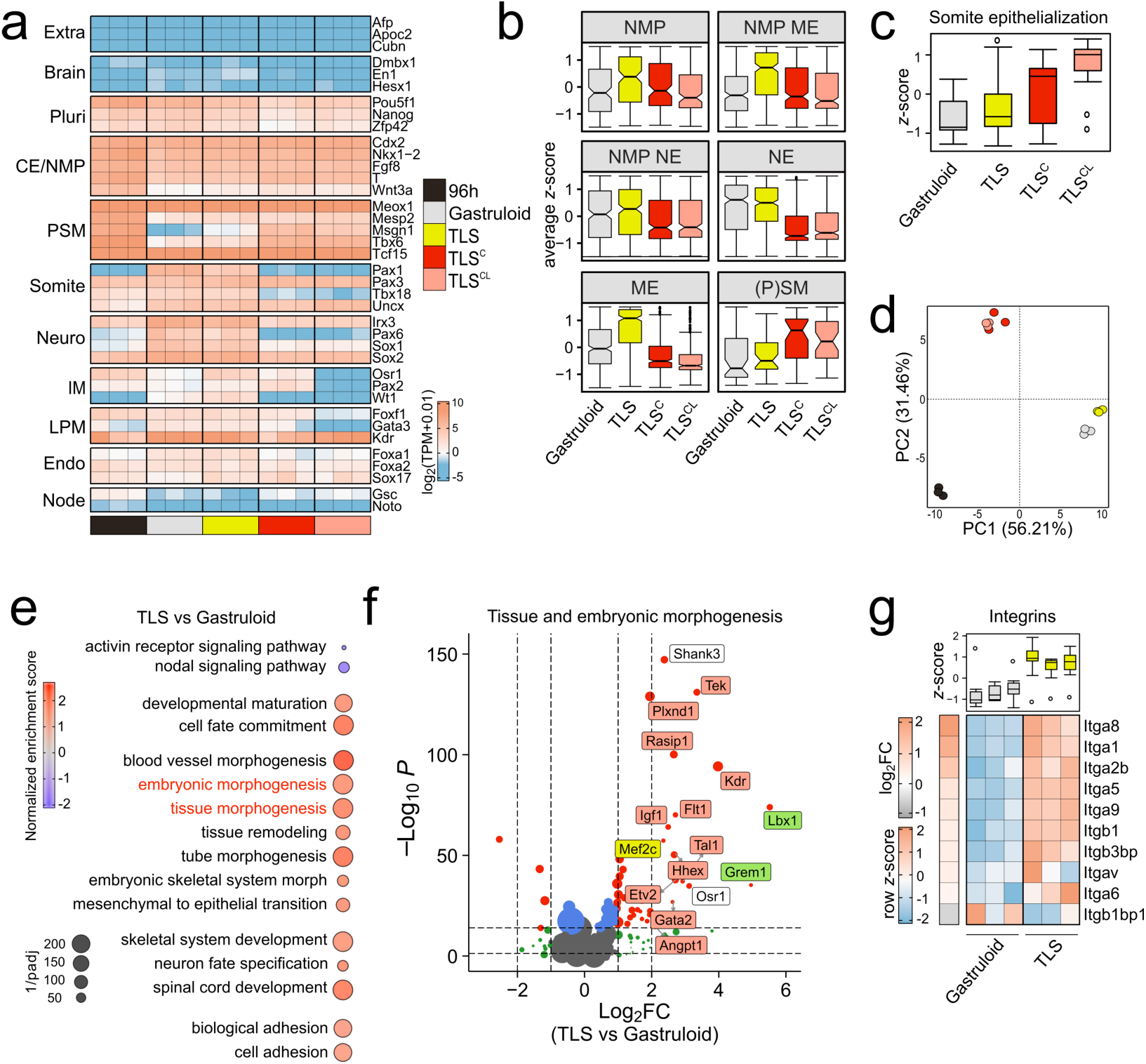
Gene expression differences between gastruloids and TLS models. **a**, Heatmap of log_2_(TPM+0.01) expression (TPM, Transcripts Per Million) of selected genes associated with development of indicated embryonic structures in 96h aggregates, 120h gastruloids, 120h TLS, 120h TLS^C^ and 120h TLS^CL^, as measured by RNA-seq. Replicates were derived from pools of independent biological samples (see **Extended Data** Fig. 3a for exact experimental set-up). CE, caudal end, NMP, neuromesodermal progenitors, PSM, presomitic mesoderm, LPM, lateral plate mesoderm, IM, intermediate mesoderm. **b**, Box plots showing distribution of marker genes for indicated cell types. Boxes indicate interquartile range. End of whiskers represent minimum and maximum. Black dots indicate outliers. Notches are centered on the median. List of genes used for each category in **Supplmentary Table 1. c**, Boxplot like in **b** representing average *z*-score per column (pool of 3 replicates) for somite epithelialization factors (**Extended Data** Fig. 3f for genes). **d,** PCA analysis of samples from **a** with color coding of individual samples (dots) as in **a**. PC1 and PC2 represents the two components with highest percentage of explained variance. **e,** Selected significant terms of Gene Set Enrichment Analysis (GSEA) enriched in TLS as compared to gastruloids of 120h. Full list of significant (FDR<0.05) terms is provided in **Supplementary Table 2**. **f,** Volcano plot of genes involved in tissue and embryonic morphogenesis. Dot size scales with log_2_ of absolute expression. Red dots, absolute log_2_FC>1 and padj<10e^-15^. Green dots, absolute log_2_FC>1. Upper dotted line, padj (FDR)=10e^-15^; bottom dotted line, padj (FDR)=0.05. Green label, involved in somitogenesis; orange label, involved in blood vessel development; yellow label, involved in both^40–48^. **g,** Heatmap of scaled expression (row *z*-score) of integrins with significantly different expression (padj(FDR)<0.05) in gastruloids vs 120h TLS. Boxplot represents *z*-score per column (sample), with boxes indicating interquartile range, end of whiskers representing minimum and maximum, dots showing outliers and central line representing median. Every column represent one of three biological replicates.

To characterize our structures in more detail we performed RNA-seq analysis (**Extended Data Fig. 3a**), and found that TLS model the post-occipital embryo, similar to gastruloids, based on selected marker genes (**Fig. 2a; Extended Data Fig. 3b**)^3^. Compared to TLS, both TLS^C^ and TLS^CL^ showed a significant upregulation of genes involved in (pre)somitic development (e.g. *Tbx6*, *Msgn1*, *Hes7*^7, 8, 11^) at the expense of NE marker genes (e.g. *Sox1*, *Pax6*, *Irx3*^11–13)^, corroborating the flow cytometry and imaging results (**Fig. 2a; Extended Data Fig. 3b-d**). The analysis of marker gene sets for NMPs, their direct descendants undergoing lineage choice (NMP ME & NMP NE), and for committed NE and ME cells substantiated this finding^11^. Compared to TLS, TLS^C^ and TLS^CL^ displayed reduced expression of markers in all clusters including ME (**Fig. 2b**). (P)SM specific markers, however, were on average upregulated, whereas intermediate ME (IM) and lateral plate ME (LPM) markers were downregulated in TLS^C^, and further reduced in TLS^CL^ (**Fig. 2b; Extended Data Fig. 3c,e**)^3, 8^. These data are in line with the known role of WNT-versus BMP-signaling in conferring PSM versus IM and LPM subtype identity (**Fig. 2a; Extended Data Fig. 3e**)^7^.

**Fig. 3.**
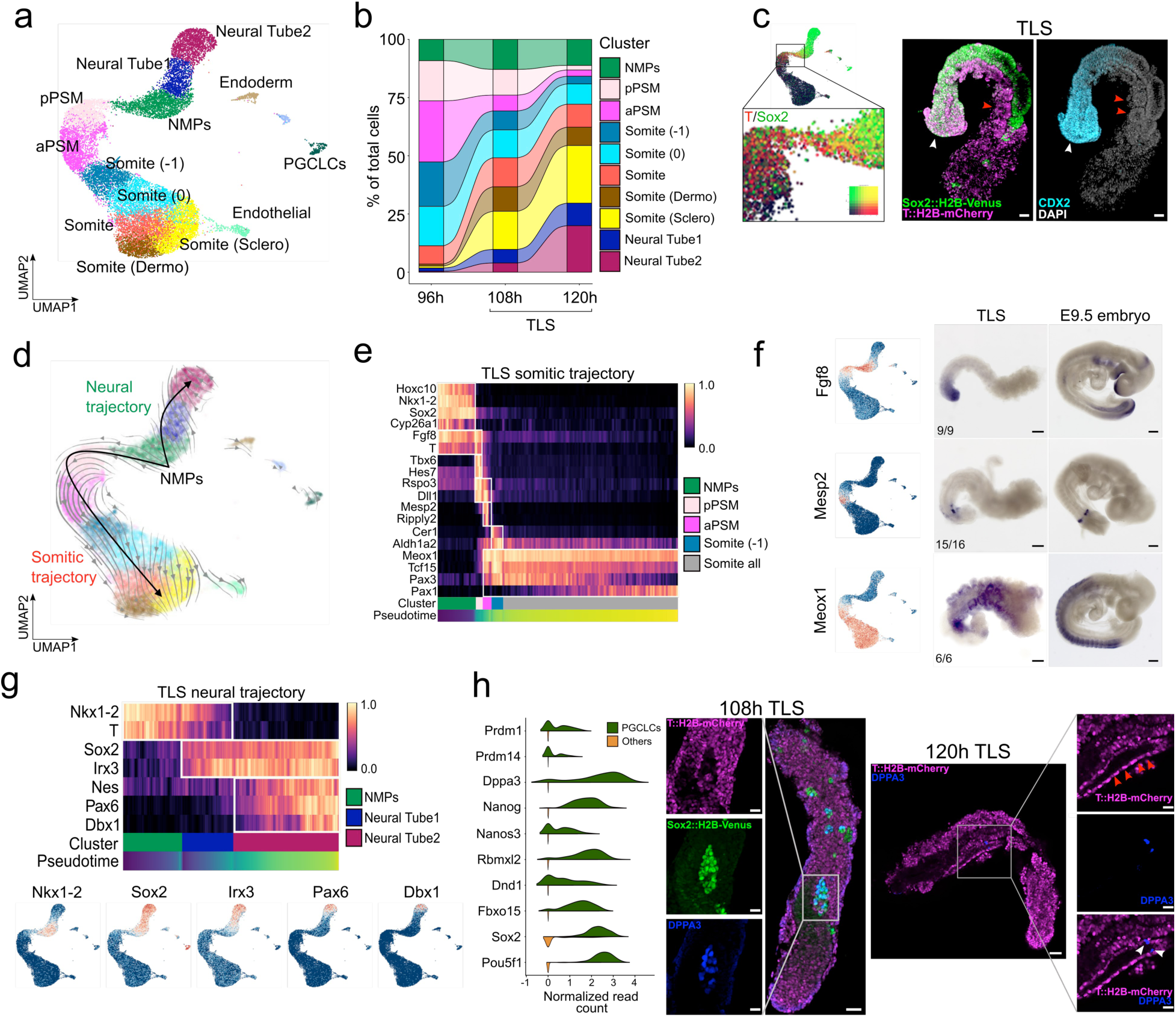
Single-cell RNA-Sequencing of TLS. In total, 20,294 cells were sampled from 96, 108, and 120 hour TLS (see **Extended Data Fig. 5a** for experimental set-up). **a,** UMAP (Uniform Manifold Approximation and Projection) coloured by the fourteen clusters identified. **b**, Alluvial plot of percentage of neuromesodermal progenitors (NMPs), posterior presomitic mesoderm (pPSM), anterior PSM, somitic and neural tube cells over time. **c,** NMPs co-express Sox2 and T (left panel, blending expression with blend.threshold=0.4), and reside at the posterior end of TLS (confocal sections, right panel; white arrowheads, NMPs; red arrowheads, somites). Scale bars 50μm. **d**, UMAP coloured by identified clusters with trajectories inferred from RNA Velocity. Grey arrow flows represent calculated velocity trajectories. **e**, Heatmap with scaled expression of genes involved in somitogenesis in 9004 cells from 120h TLS^MG^ rooted in NMPs and ordered by pseudotime. **f**, UMAP coloured by expression of indicated genes (left panel), and whole-mount *in situ* hybridization for the same genes in TLS and E9.5 embryos (right panel). Numbers indicate the fraction of TLS with embryo-like expression. Scale bars TLS 100μm, embryo 200μm. **g**, Heatmap with scaled expression of genes involved in neural development in 9004 cells from 120h TLS rooted in NMPs and ordered by pseudotime (upper panel) and UMAP coloured by expression of indicated genes (bottom panel). **h**, Split violin plots of expression of marker genes for primordial germ cell like cells (PGCLC, left panel), and confocal section of TLS showing SOX2^VE-high^/DPPA3^+^ PGCLCs at 108h, and DPPA3^+^ cells in close contact with the T^mCH+^ gut-like-structure. Scale bars 50μm for 108h left panel and 120h overview, 25μm for 120h magnifications. Red arrowheads indicate gut-like-structure, white arrowheads indicate DPPA3^+^ PGCLCs.

We next searched for gene expression differences that might underlie the improved physical separation of somites observed in TLS^C^ and TLS^CL^. Among the most strongly upregulated genes compared to TLS was *Wnt6*, which acts as a somite epithelialization factor *in vivo* (**Extended Data Fig. 3f**)^14^. In addition, multiple ephrins and their receptors, and other factors involved in somite ephithelialization were upregulated (**Fig. 2c; Extended Data Fig. 3f**)^15, 16^. Expression changes of selected somite polarity markers was observed concomitantly with changes in their inducers, in agreement with the role that WNTs, in concert with SHH, BMPs and their antagonists play in somite compartmentalization *in vivo* (**Extended Data Fig. 3g,h**)^7, 17^. Thus, exposure to CHIR or CHIR/LDN improved segment boundary formation, but affected somite patterning.

Principal Component Analysis (PCA) indicates a high transcriptional similarity between gastruloids and TLS despite profound morphological differences (**Fig. 2d**). The latter are better highlighted by Gene-Set-Enrichment-Analysis (GSEA), which shows that matrigel embedding promotes tissue morphogenesis and remodeling (**Fig. 2e; Extended Data Fig. 3i**). Zooming in on embryonic and tissue morphogenesis gene sets showed that upregulated genes also comprise markers of blood vessel development suggesting the induction of capillary morphogenesis in TLS (**Fig. 2f; Extended Data Fig. 3j**). Cell-cell and cell-matrix interactions play pivotal roles in morphogenesis, with important functions for (proto)cadherins, ephrins and integrins in tissue formation and organogenesis^18–23^. GSEA indeed revealed an enrichment of cell adhesion terms and overall a significant upregulation of corresponding marker genes in TLS (**Fig. 2e; Extended Data Fig. 4a,b**). The most pronounced increase was observed for integrins, transmembrane receptors mediating cell adhesion to the ECM important for e.g. neural tube formation, blood vessel development, and segmentation (**Fig. 2g; Extended Data Fig. 4a,b**)^20, 21, 24–26^. Of note, integrin ligands important for somite boundary formation (fibronectin, collagen IV, laminin alpha1 and laminin alpha5) were expressed at the same levels in gastruloids (**Extended Data Fig. 4c**)^24, 25, 27, 28^. Taken together, our molecular data confirm activation of morphogenetic programs through matrigel embedding that may be linked to the upregulation of cell adhesion molecules.

**Fig. 4.**
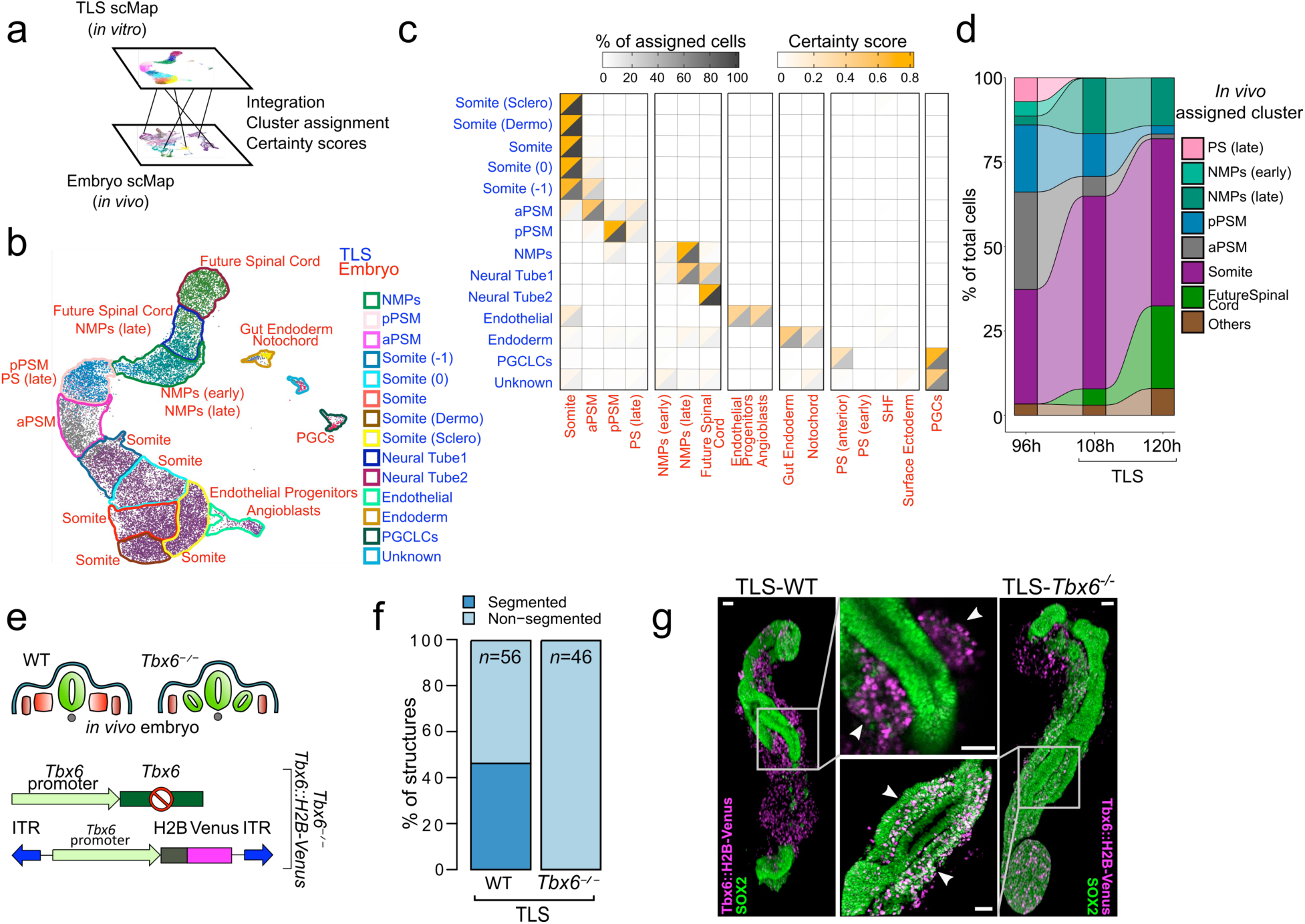
TLS cell types are embryo-like and recapitulate the *Tbx6^-/-^* phenotype at the molecular and morphological level. **a**, Comparative transcriptome analysis of TLS and post-occipital E7.5 and E8.5 embryo at the single-cell level. **b**, TLS UMAP coloured by assigned embryonic cell type. TLS clusters are projected as corresponding coloured contours. Blue font, TLS clusters; Red font, embryo clusters. **c**, Split heatmap with percentage of assigned cells (dark grey) and certainty score (orange) for TLS cells from the indicated cluster upon unbiased mapping to the *in vivo* counterpart. Font colour code as in **b**. **d**, Alluvial plot of percentage of cells assigned to indicated *in vivo* cluster in 96, 108 and 120h TLS. **e,** Schematic of *Tbx6^-/-^* phenotype *in vivo* (upper panel) and schematic view of *Tbx6::H2B-Venus; Tbx6^-/-^* mESC derivation. **f,** Quantification of segmentation phenotype in TLS-*Tbx6^-/-^*. Data represent 3 different experiments performed with 2 independent mESC lines of each genotype. **g,** Formation of ectopic neural tubes in TLS-*Tbx6^-/-^*. Green, SOX2; Magenta, Tbx6^VE^. White arrowheads indicate Tbx6^VE+^ somites in WT, and Tbx6^VE+^/SOX2^+^ ectopic neural tubes in *Tbx6^-/-^*. Scale bars 50μm.

Based on the above results we decided to focus on the TLS condition for an in-depth characterization, as it produced the most *in-vivo*-like configuration. First, to ensure reproducibility^29^, we evaluated variability between nine independent structures using the quantitative expression of a panel of 41 developmental genes. The data demonstrated high correlation between TLSs and revealed that they most closely resemble the E8.5 post-occipital stage (**Extended Data Fig. 5a**). The analysis of individual genes showed that endothelial, NMP, NE, and somitic ME genes were reproducibly expressed at similar levels across all replicates, whereas higher variation was observed in the expression of PSM, endodermal, and pluripotency genes (**Extended Data Fig. 5b,c**).

Next we performed scRNA-seq analysis on a total of 20,294 post-processed cells sampled from TLS at 96, 108, and 120 hours (**Extended Data Fig. 6a**). Clustering analysis revealed 14 different cell states with the larger ones corresponding to derivatives of the PSM and NE that flank putative NMPs. Smaller clusters comprised endoderm, endothelial cells and Primordial Germ Cell Like Cells (PGCLCs) (**Extended Data Fig. 6b**). The main clusters organized into a continuum of states recapitulating spatio-temporal features of the developing post-occipital embryo (**Fig. 3a**). Across the three time points sampled progenitor subtypes gradually decreased in favor of more mature neural and somitic cells as development progresses (**Fig. 3b; Extended Data Fig. 6c**). As expected, putative NMPs co-expressed T^mCH^, Sox2^VE^ and CDX2, and showed an *in-vivo*-like NMP signature and location (**Fig. 3c; Extended Data Fig. 6d**)^10, 11, 30^. RNA Velocity analysis revealed neural and somitic trajectories rooted in the NMPs, suggesting that TLS development recapitulates the developmental dynamics observed in the post-occipital mid-gestational embryo (**Fig. 3d; Extended Data Fig. 7a**)^31^. *In vivo*, NMPs and their PSM descendants are arranged in an order of progressive maturity along the posterior to anterior axis^7^. In line with this, ordering of cells along a pseudo-temporal trajectory showed that the somitic trajectory reflects the genetic cascade observed in the embryo (**Fig. 3e; Extended Data Fig. 8a**). For example, the trajectory from Fgf8^+^ NMPs and PSM via the determination front marked by

Mesp2 to Meox1^+^ somites was faithfully recapitulated and the embryo-like spatial arrangement was confirmed by whole-mount *in situ* hybridization (**Fig. 3f**)^7^. Likewise, the genetic cascade from NMPs to neural progenitors in TLS reflected its *in vivo* counterpart in space and time (**Fig. 3g**). Notably, subclustering of the neural cells demonstrated that TLS generate both dorsal and ventral neural subtypes, with dorsal subtypes being more prevalent (**Extended Data Fig. 8b**)^32^. The analysis of *Hox* gene expression at consecutive time points demonstrated *in-vivo*-like collinearity, as described for gastruloids (**Extended Data Fig. 8c**)^3^. To test if TLS somites establish dorsal-ventral (DV) and anterior-posterior (AP) domains, we reclustered all somitic cells. At 96h we detected two main groups, corresponding to the Uncx^+^ posterior and Tbx18^+^ anterior somite domains, in line with AP compartmentalization established during segmentation (**Extended Data Fig. 9a,b**)^7^. At 120h, we found distinct clusters of Pax3^+^ dorsal and Pax1^+^ ventral cells, and a small cluster of Lbx1^+^/Met^+^ putative migratory muscle precursors (**Extended Data Fig. 9c-f**)^7, 33^. In addition, Scx^+^ syndetome cells were detected (**Extended Data Fig. 9g**), and Uncx and Tbx18 expression were anti-correlated (**Extended Data Fig. 9h**). An unexpected finding was the identification of PGCLCs, since they have not been observed prior in gastruloids^3^. In the embryo, nascent PGCs can be identified at E7.5 as a group of DPPA3^+^ cells, which are generated in the posterior primitive streak and later migrate along the hindgut to the gonads^34^. We assigned PGCLC identity using characteristic PGC genes and identified their location in our structures (**Fig. 3h; Extended Data Fig. 10**). At 108h we find a group of Sox2^VE-high^ cells that co-expressed DPPA3 (**Fig. 3h; Extended Data Fig. 10a**). At 120h, Sox2^VE-high^ cells were detected in contact with FOXA2^+^ cells, and DPPA3^+^ cells in contact with a T^mCH+^ gut-like epithelial structure (**Fig. 3h; Extended Data Fig. 10b**). These data show that trunk-like structures indeed contain cells displaying characteristics typical for PGCs. To investigate how close the cellular states identified in TLS resemble those in embryos, we mapped our single-cell transcriptomes to a scRNA-seq compendium of post-occipital embryonic cellular subtypes (**Fig. 4a**)^35^. The data revealed globally a high accordance of TLS and embryonic cell types including characteristic marker genes and pairwise comparison of mapped clusters identified only a small fraction of differentially expressed genes (**Fig. 4b,c; Extended Data Fig. 11a-d**). Of note, PS- and early-NMP-like cells are exclusively present at 96h and replaced by late-NMP-like cells at 108h and 120h (**Fig. 4d**). Taken together, our scRNA-seq analyses demonstrate that the trunk-like structures execute gene-regulatory programs in a spatiotemporal order that resembles the embryo.

Finally, to explore the utility of trunk-like structures further, we next tested if they could reproduce an embryonic mutant phenotype caused by gene ablation in a proof of concept experiment. *In vivo*, loss of *Tbx6* results in an expansion of NE and the formation of ectopic neural tubes at the expense of PSM and somites (**Fig. 4e**)^36^. We deleted *Tbx6* from *Tbx6::H2B-Venus* (*Tbx6*^Ve^) mESC, generated TLS and found a clear failure to form somites, which could also not be rescued by CHIR or CHIR/LDN (**Fig. 4e,f; Extended Data Fig. 12a**). Quantitative PCR analysis on FACS-purified Tbx6^VE+^ cells revealed upregulation of neural markers at the expense of (P)SM markers in *Tbx6^-/-^* cells, thus recapitulating the *in vivo* phenotype at the molecular level (**Extended Data Fig. 12b**). Finally, whole-mount immunofluorescence analysis for SOX2 showed that TLS-*Tbx6^-/-^* generated ectopic Tbx6^VE+^ neural tubes, whereas TLS^C^ and TLS^CL^ formed an excess of morphologically indistinct SOX2^+^ tissue (**Fig. 4g; Extended Data Fig. 12c,d**).

Here we report the generation of trunk-like structures and demonstrate that they faithfully reproduce important features of post-occipital embryogenesis, as confirmed by morphogenetic and transcriptional criteria. Importantly, to our best knowledge axial elongation together with neural tube, gut and somite formation in combination with the generation of PGCLCs, solely by *in vitro* culture of stem cells, has been achieved for the first time. TLS therefore provide a powerful *in vitro* system for investigating the dynamics of embryo patterning and morphogenesis at the single cell level and in molecular detail, allowing to “understand the whole from the parts”^1^. Towards this latter goal, comparative analysis of gastruloids and TLS will help to understand how cell-cell and cell-matrix interactions control embryonic architectures^1^. In this regard, cell type specific expression of cell adhesion molecules revealed by our single-cell TLS atlas will provide guidance for future work (**Fig. S13**). Although TLS display the closest *in-vivo*-like configuration, we envision that our CHIR and CHIR/LDN models can become important systems for studying the morphogenetic aspects of somitogenesis in high-throughput. Our TLS already display remarkable reproducibility both at the morphological and molecular level. Nevertheless, we realize the importance of further integrated comparative analysis of single TLS at single-cell resolution, such as recently published for brain and kidney organoids^37, 38^. Finally, *in vivo* somites are formed in a rhythmic process involving an oscillator - the segmentation clock^39^. We have observed that somite formation in TLS can occur sequentially at an embryo-like pace, and specific control genes involved in the segmentation process are correctly expressed in aPSM as expected, suggesting that the clock also ticks in the PSM of TLS (**Extended Data Fig. 14; Supplementary Video 1**).

In sum, our data demonstrate that the morphogenetic potential of mESC-derived aggregates is unlocked by providing an ECM surrogate. The resulting trunk-like structures provide a scalable, tractable, readily accessible platform for investigating lineage decisions and morphogenetic processes shaping the mid-gestational embryo at an unprecedented spatiotemporal resolution.

## Acknowledgments

We are grateful for the support and feedback received from members of the Herrmann & Meissner laboratories, in particular Stefanie Grosswendt and Atsuhiro Taguchi. We thank Dijana Micic & Judith Fiedler for animal care, Norbert Mages for assistance with (sc)RNA-Seq, Claudi Giesecke-Thiel & Uta Marchfelder for assistance with FACS, Thorsten Mielke & Beatrix Fauler for help with microscopy, Fabian Tobor and Polly Burton for technical assistance, and Nikolaus Rajewsky (MDC/BIMSB) for providing access to the NanoString. **Funding:** J.V.V. was partly funded by an Alexander von Humboldt Fellowship. This work was supported by NIH grant HG006193 (A.M.) and the Max Planck Society.

## Author contributions

B.G.H. initiated the study; J.V.V. and B.G.H. conceived the project; J.V.V., A.M. and B.G.H. supervised the project. J.V.V., A.B. and L.H. designed, performed and quantified most experiments. H.K. performed bulk & sc-RNA-Seq computational analysis with help of J.V.V. and A.B. M.S-W., D.S., F.K. and M.P. generated mESC reporter lines. L.W. performed tetraploid complementation. S.H. performed pilot experiments to optimize culture media. R.B. helped with image acquisition and analysis. B.T. supervised next-generation sequencing. J.V.V. drafted the first version of the manuscript. The final manuscript was written by J.V.V, A.B., H.K., A.M. and B.G.H.

## Data and materials availability

All data is available in the main text or the supplementary materials. Bulk and single-cell RNA-Seq data have been deposited in the Gene Expression Omnibus (GEO) under accession code GSE141175. All computational code used in this study is available upon request.

## Methods

### Animal work

All animal work was approved by the local authorities (LAGeSo Berlin, license numbers G0247/13 and G0243/18). *T::H2B-mCherry/Sox2::H2B-Venus* embryos were generated via tetraploid complementation^49^. For embryo isolation, mice were sacrificed by cervical dislocation and uteri were dissected in PBS.

### Mouse embryonic stem cell culture

All medium compositions are listed in **Supplementary Table 3**. All mouse embryonic stem cells (mESCs) used in this study were male and from an F1G4 background^50^. mESCs were maintained on 6cm plates (Corning 430166) gelatinized with 0.1% gelatin (1:20 dilution of 2% gelatin (*Sigma G1393*) in tissue-culture grade H_2_O) and coated with mitotically inactive primary mouse embryo fibroblasts (3-4×10^4^cells/cm²) with standard mESC medium containing 15% FCS and 1000 U/ml leukemia inhibitory factor (*LIF, Chemicon ESG1107*) at 37°C and 5% or 7.5% CO_2_. mESCs were split every second day with a dilution suitable to the proliferation velocity (between 1:5 and 1:9). ES+LIF medium was refreshed daily. For splitting, media was aspirated and cells were washed once with PBS and trypsinized (Tryspin-EDTA (0.05%) (Gibco 25300054)) for 5-10’ at 37°C. Trypsin was neutralized by 3ml ES+LIF and cells centrifuged for 5’ at 1000 rpm, after which the pellet was resuspended in ES+LIF. For freezing of mESCs, cell pellets were resuspended in ES medium with 20% FCS, and mixed in a 1:1 ratio with ES freezing medium. Cells were frozen down o/n in the –80 °C and transferred to liquid nitrogen the next day.

### Generation of transgenic mouse embryonic cell lines

The mouse embryonic cell lines used for the experiments were all derived from F1G4 cells. To generate the fluorescent reporter constructs, we genome-engineered the mouse BACs *RP24-530D23* (*T*), *RP23-249O15* (*Sox2*) and *RP23-421P23* (*Tbx6*), containing ∼180 −230kb of the C57BL/6 mouse genome surrounding the respective loci, via Red/ET recombineering^51^. In short, the starting codon (ATG) of the genes was replaced with a reporter gene containing H2B-mCherry (for *T*) or H2B-Venus (for *Sox2*/*Tbx6*), followed by the rabbit *b-globin* polyadenlylation signal and an *FRT*-flanked selection cassette (hygromycin or neomycin), driven by the *Pgk* promotor. For random integration, 5μg of the modified BACs was linearized with PI-SceI (NEB) and electroporated into 3×10^6^ mESCs. Selection was performed using mESC medium containing 250μg/ml neomycin and 150μg/ml hygromycin for the T::H2B-mCherry;Sox2:H2B::Venus reporter line. For the Tbx6::H2B-Venus reporter line 350μg/ml hygromycin was used for selection. After selection, single clones were picked, expanded and checked for BAC integration by PCR. Two independent *Tbx6::H2B-Venus* mESC lines were generated. In both lines, the endogenous *Tbx6* locus was targeted by CRISPR (**Supplementary Table 4**) to create null mutants. To this end, a double nicking approach with four guide RNAs^52^ was used. The specific target sequences contained 5’-*GN_19_NGG*-3’’, with *N* being any arbitrary nucleotide, *G* being guanine and *NGG* the Protospacer adjacent motif and *GN_19_* the guide RNA. If there was no guanine at the 5’-end of the template sequence, an extra guanine had to be added 5’ to the other 20 bases. Single stranded oligonucleotides (**Supplementary Table 4**) with an added 5’-*CACC-3’* at 5’-end and 5’-*AAAC*-3’ at 5’-end of the complementary strand were annealed in 10xT4 ligation buffer *(Promega M1801)* by continuous cooling from 95°C to 25°C. Annealed oligos were cloned into the *BbsI* site of px335A_hCas9_D10A_G2P (gift from Dr. Boris Greber) (**Supplementary Table 4**) containing expression cassettes for hCas9 nickase, guide RNA and puromycin resitance.

One day prior to transfection, 3×10^5^ mESCs cells were seeded on fibroblast coated 6-well plates *(Costar 3516)* with 3 ml ES+LIF per well. After overnight incubation, the medium was refreshed in the morning. Mixes of 110µl Opti-MEM reduced serum medium *(Gibco 31985062)* plus 25µl Lipofectamine2000 *(Thermo Fisher 2125239)* and 125µl Opti-MEM plus 8µg per vector were prepared. 125µl of each mix were combined and incubated for 15 min at room temperature before being transferred to 1.25ml ES+LIF without Penicillin/streptomycin. After 5h of incubation with the transfection mix, ESCs were split and plated on puromycin resistant feeders. 24h after transfection, transient selection was started with ES+LIF containing 8 µg/ml puromycin *(Gibco 10130127)* at day 1 and day 2 and ES+LIF containing 4 µg/ml puromycin at day 3. After approximately one week, single clones were picked and expanded on 96-well plates *(Costar 3596)*. Clones were genotyped by PCR using primers spanning Exons 1-4. Deletion breakpoints were analyzed by Sanger sequencing of purified PCR fragments.

### Generation of gastruloids

All medium compositions are listed in **Supplementary Table 3**. Gastruloids were generated as described previously^53^, with some minor modifications. First, mESCs were feeder freed. To this end, mESCs were trypsinized on the feeder plate as described above, washed with ES+LIF and resuspended in 2ml ES+LIF. On three gelatinized (0.1% gelatin) wells of a 6-well plate, cells were sequentially plated for 25’, 20’ and 15’ during which cells were kept in the incubator at 37°C and 5% or 7.5% CO_2_. With each transfer, cells were triturated to maintain a single cell suspension. Feeder-freed mESCs were then washed once in 5 ml PBS containing MgCl_2_ and NaCl (Sigma A8412) and once in 5 ml NDiff227 (*Takara Y40002*). mESCs were then pelleted by centrifugation for 5’ at 1000rpm and resuspended in 500µl of NDiff227. 10µl of the cell suspension was mixed with 10 µl of Trypanblue (*Bio-Rad 1450021*) for automated cell counting with *Luna Automated Cell Counter*. 200-250 live cells were then plated in a volume of 30 to 40µl NDiff227 into each well of a 96-well round bottom, low attachment plate (*Cellstar 96 well suspension culture plate (655185)* or *Costar 7007 ultra-low attachment 96 well plate (7007)*). Cells were then allowed to aggregate for 48 h. After these 48h cells were pulsed with 3µM CHIR99021 (CHIR, *Merck Millipore*) in 150µl NDiff227. Between 72 and 120h, medium was refreshed every 24h by removing 150µl of the old media and adding the same volume of new, pre-incubated (37°C and 5% or 7.5% CO_2_) NDiff227. For gastruloids treated with CHIR and CHIR+LDN, 5µM CHIR with or without 600nM LDN-193189 (LDN, *Reprocell*) was added from 96h to 120h. For controls, an equal volume of diluent (DMSO) was added.

### Generation of trunk-like-structures

All medium compositions are listed in Data S3. The gastruloid protocol described above was followed until 96h. Gastruloids were then embedded in 5% Growth-Factor-Reduced Matrigel (MG) (*Corning 356231*). To this end, fresh NDiff227 medium was pre-incubated for at least 20’ at 37°C and 5% or 7.5% CO_2._ Pre-incubated medium was then put on ice for 5’, after which MG was added to achieve a final concentration of 5% in the culture wells. Medium was then put on room temperature for 5’, during which 150µl of old medium was removed from the aggregates. New medium with MG (150µl) was then added, and the cultures were returned to the incubator and further cultured at 37°C and 5% or 7.5%. TLS cultures were allowed to settle for at least 30’ before proceeding to further experimentation (e.g. live imaging). For TLS treated with CHIR and CHIR+LDN, 5µM CHIR with or without 600nM LDN was added from 96h to 120h prior to adding the MG. For controls, an equal volume of diluent (DMSO) was added.

### Whole-mount Immunofluorescence

Collected embryos were washed twice in PBS and then fixed in 4% PFA for 30’ under rotation at 4°C, washed 3x with PBS, and stored in PBS until immunofluorescent staining was performed. Gastruloids or trunk-like-structures were picked using a p200 pipette with the tip cut-off at the 50µl mark. Gastruloids were washed twice with PBS and then fixed in 4% PFA for 1 h in glass vials (Wheaton 224882) at 4°C on a roller. For trunk-like-structures, individual structures were picked using a p200 pipette with the tip cut-off at the 50µl mark, and transferred to either 96-well plates (Costar 3596) or Ibidi 8-well glass-bottom plates (*Ibidi 80827*). Trunk-like-structures were washed twice with PBS + MgCl_2_ and NaCl + 0.5% BSA (Sigma A8412), once with PBS, and then fixed in 4% PFA for 1hr at 4°C on a rocking platform. Subsequently, gastruloids or trunk-like-structures were washed twice in PBS for 5 min, permeabilized by 3 x 20’ incubation in 0.5% Triton-X/PBS (PBST) and blocked in 5% fetal calf serum/PBST (blocking solution) overnight at 4°C. For antibody staining, gastruloids/trunk-like-structures were transferred to Ibidi 8-well glass bottom plates. Primary antibody incubation was performed in blocking solution for 48-72h at 4°C, after which gastruloids/trunk-like-structures were washed three times with blocking solution and three times with PBST. After the last washing step, Gastruloids/trunk-like-structures were incubated in blocking solution o/n at 4°C. The next day, secondary antibodies diluted in blocking solution were added, and structures were incubated for 24h at 4°C. Afterwards, gastruloids/trunk-like-structures were washed three times with blocking solution and three times with PBST. The last PBST washing step after secondary antibody incubation included DAPI (0.02%, Roche Diagnostics 10236276001). DAPI was incubated o/n and washed off once with PBST. All primary and secondary antibodies are listed in **Supplementary Table 3**.

### Tissue clearing

Prior to imaging, embryos, gastruloids and trunk-like-structures were cleared with RIMS (refractive index matching solution). To this end, samples were washed twice with PBS for 10 min, post-fixed in 4% PFA for 20 minutes and washed three times with 0.1M phosphate buffer (PB, 0.025M NaH_2_PO_4_ and 0.075M Na_2_HPO_4_, pH 7.4). Clearing was performed by incubation in RIMS (133% w/v Histodenz (Sigma-Aldrich D2158) in 0.02M PB) on a rocking platform at 4°C for at least one to several days.

### Imaging

Gastruloids and trunk-like-structures stained with antibodies or carrying fluorescent reporters were imaged with the Zeiss AxioZoom v16 (wide-field), Celldiscoverer7 (wide-field), Zeiss LSM710 (laser-scanning microscope with Airyscan) or Zeiss LSM880 (laser-scanning microscope with Airyscan) with appropriate filters for mCherry, Venus, DAPI, Alexa Fluor 488, Alexa Fluor 568, Alexa Fluor 647, and combinations thereof. Embryos were imaged with Zeiss Lightsheet Z1 with appropriate filters for mCherry and Venus. Whole-mount in situ hybridization images were acquired with Zeiss AxioZoom v16. Post-acquisition image processing was performed using Zen and/or Arrivis. For live imaging of TLS in 96-well plates we used the Celldiscoverer 7 (Zeiss) with incubator chamber temperature set at 37°C and CO2 content at 5%. Acquisition intervals ranged from 15-30 minutes.

### Morphometric analysis of gastruloids and trunk-like-structures

For quantification of the number of structures with a T^+^-pole, we employed the *T::H2B-mCherry* reporter line. If a structure displayed multiple axes of elongation, it was sufficient for one end to have a *T::H2B-mCherry*^+^ pole to be scored as having a “T-pole”. We defined gastruloids/trunk-like-structures as “segmented” when at least four neighboring segments had developed along the anteroposterior axis at 120h. Finally if a structure displayed two or more axes of elongation, it was scored as “multiple axes”, if showing just one as “one axes”, if none “no elongation”. For all conditions, three independent experiments, each one including each treatment, were analyzed. Structures that grew out of focus in a way that they could not be rated for one of the categories were excluded for the respective category.

### Flow cytometry and FACS

Individual trunk-like-structures were washed with PBS+0.5%BSA 2 times, after which 50μl of trypsin was added. Trunk-like-structures were then incubated for 10’ at room temperature, after which samples were dissociated by trituration for 50 times using a p200 pipette to achieve a homogenous suspension. The reaction was then stopped with the addition of 100μl of 5% BSA in PBS. Before cell counting and/or sorting on a FACS Aria II (Becton Dickinson) or a BD FACSCelesta Flow Cytometer (Becton Dickinson, counting only) the cell suspension was filtered through a 35μm mesh. Equal numbers of samples for each condition were harvested in three independent experiments. For FACS, samples were dissociated as described above, and the Tbx6^VE+^ and Tbx6^VE-^ fraction was sorted in 1.5ml low-binding tubes with 350μl RLT (Qiagen) + 1% v/v B-mercaptoethanol (Sigma). Flow cytometry data were later analyzed using FlowJoV10.

### RNA isolation, reverse transcription and quantitative PCR

RNA isolation was performed using the RNeasy Micro Kit (Sigma 74004) according to the manufacturer’s instructions with the following modifications: genomic DNA was digested on column with the addition of an extra 1μl of RNase-free DNase I (Roche) to ensure complete digestion, and after the 80% ethanol column wash and column centrifugation the remaining ethanol was removed with a p10 pipette tip and columns were left to air-dry for 5’. Quantitative reverse transcriptase PCR (qRT-PCR) was then performed using a two-step protocol. First, RNA was reverse transcribed using the Quantitect Reverse Transcription Kit (*Qiagen*), according to the manufacturer’s instructions. qPCR was carried out on a StepOnePlus Real-Time PCR System (*Life Technologies*) using GoTaq qPCR Master Mix (*Promega*) with validated gene-specific primers (**Supplementary Table 4**). Fold change was calculated from ΔCt using Tbp as housekeeping gene (**Supplementary Table 5**).

### Measuring of segment size

Segment sizes were measured in Fiji 1.8.0 using the images acquired in the T^mCh^ channel. For each structure, the four newest neighboring segments were measured counting from the posterior end. In case of bilateral segments, only segments on one side were measured. For the ‘bunches of grapes’ in TLS^C^ and TLS^CL^, four segments along the antero-posterior axis were measured counting from the posterior end. Length was defined as parallel to the anterior-posterior axis, width was defined as perpendicular to the anterior-posterior axis.

### Whole-mount in situ hybridization

#### Probe synthesis

For synthesis of DIG labelled RNA probes, plasmids containing the cDNA of interest and promoters for sense and anti-sense strand synthesis were used from the MAMEP database (http://mamep.molgen.mpg.de). In order to obtain a sufficient amount of material, some plasmids were first retransformed into *E. coli* (DH5α) and afterwards isolated with a mini-prep kit (Qiagen) according to the manufacturer’s instructions as described below. To verify the identity of the vector and insert of each plasmid, plasmids were digested by restriction enzymes and loaded on a 1.0% agarose/TAE gel. Probes were synthesized by *in vitro* transcription using a PCR product of the desired cDNA or the linearized plasmid.

#### Retransformation in DH5α and mini-prep

DH5α cells were thawed on ice and 1µl of the plasmid was added to 100µl of competent cells and incubated on ice for 30’. Subsequently, cells were heat shocked at 42°C for 45” in a water bath and then immediately cooled down on ice. 500µl of LB medium was added and the mix was incubated for 1h at 37°C (heating block) under shaking. Then, 25µl and 250µl of the mixture were plated on separate Agar-plates containing Ampicillin (Amp) for selection of transformed bacteria (all vectors carried a gene for Ampicillin resistance). The plates were incubated overnight at 37°C. The next day, single colonies were picked in 5ml LB+Amp and incubated overnight at 37°C under shaking. Plasmids were isolated with the QIAprep Spin Miniprep Kit (*Qiagen*) following the instructions of the Quick-Start Protocol using a centrifuge for processing. DNA concentration was measured with a Nanophotometer (*Implen*).

#### Restriction Digest

After plasmid digestion (250ng of the plasmid, 2µl 10x buffer, 0.2µl 100x BSA and 2µl of each restriction enzyme in a 20µl total volume), expected band sized for the vectors and inserts was confirmed on a 1% Agarose/TAE-gel, stained with SybrSafe (1:20.000).

#### Polymerase-Chain-Reaction (PCR)

For PCR, 5µl Plasmid (1 ng/µl) was used in a 50µl total reaction volume containing 5µl 10x PCR buffer (*Invitrogen*), 1.5µl MgCl_2_ (*Invitrogen*), 10µl dNTP Mix (10 mM each nucleotide), 0.25µl forward Primer (100 µM), 0.25µl reverse Primer (100 µM) and 0.2µl Taq DNA Polymerase (5u/µl) (*Invitrogen*). For the plasmid of Uncx4.1 a different PCR strategy was used, because of its CG richness. Here, the 50µl reaction contained 5µl Plasmid (1ng/µl), 1µl dNTPS, 0.25µl U5, 0.25µl L2, 0.2µl Taq DNA Polymerase (5u/µl) (*Qiagen*) and 10µl 5x Q-solution (*Qiagen*). Primers used for the respective vectors are listed in **Supplementary Table 4**. All PCR products were checked on a 1 % Agarose/TAE-gel.

#### Linearization of plasmids for in vitro transcription

Plasmids were linearized with a restriction enzyme (1.5µg DNA, 2.5µl 10x buffer, filled up with DEPC-H_2_O to 25µl). The reaction was incubated for 1h at 37°C. Subsequently, 8µl ammonium acetate (10M) and 80µl ice-cold 100% ethanol were added, followed by centrifugation of the sample for 30’ at 4°C and 13.200 rpm. The supernatant was then removed and 150µl of 70% ice-cold ethanol was added. Sample was centrifuged again for 10’, the supernatant was removed, and the pellet was air dried and dissolved in 9.5µl DEPC-H_2_O.

#### In vitro transcription

For *in vitro* transcription, 9µl of PCR product or 1.5µg of linearized plasmid was incubated with 3µl 10x Transcription buffer, 3µl ACG nucleotides (4mM each nucleotide), 0.75µl digUTP-UTP Mix (4mM), 1.5µl DTT, 1µl RNase inhibitor and 60U of the respective RNA polymerase, in a total reaction volume (filled up with DEPC-H_2_O) of 30µl. The reaction was incubated for 2h at 37°C (T7, T3) or 40°C (Sp6). Subsequently, 3µl of RNase-free DNaseI (10u/µl) was added and the reaction was incubated for 15’ at 37°C. The RNA probe was then purified with ProbeQuant G50 Sephadex columns (*Pharmacia*). Adding 20µl DEPC-H_2_O to the probe, spinning it through the resin of the column and adding again 30µl DEPC-H_2_O increased its volume to 100µl. Aliquots of 30µl were immediately put on dry ice and stored at −80 °C. The RNA probes were checked on a 1% Agarose/TAE-gel.

#### Fixation and methanol series

Collected embryos, gastruloids and trunk-like-structures were washed twice in DEPC-PBS and then fixed in 4% PFA overnight at 4°C. The next day, samples were transferred into 100% methanol via a methanol series, including two washing steps in DEPC-PBS and a transfer from 25% to 50 % to 375% to 100% methanol/DEPC-PBS (10’ each). Upon transfer to 100% methanol, samples were washed twice in 100% methanol and stored in 100% methanol at −20°C. All steps were performed for at least 10’ at 4°C.

#### Pre-hybridization & Hybridization

Unless stated differently, all steps were performed for 10’ at 4 °C under rocking. For the composition of the solutions used for in situ (Pre-hybridization, Hybridization, Post-hybridization washes and antibody incubation, Post-antibody washes, Staining) we refer to **Supplementary Table 3**. First, samples were pre-hybridized by transferring to DEPC-PBST via a reverse methanol series (100%, 75%, 50%, 25% for 10’ each). Subsequently, samples were incubated in 6% H_2_O_2_/DEPC-PBST at 4°C, trunk-like-structures/gastruloid for 10’ and embryos for 20’, followed by three washes with DEPC-PBST. ProteinaseK/DEPC-PBST (10µg/ml) treatment was performed at 4°C, (7’ for trunk-like-structures/gastruloids, 10’ for embryos). ProteinaseK activity was quenched with Glycine/DEPC-PBST (2mg/ml) and two washes with DEPC-PBST. Subsequently, samples were post-fixed in 0.2% Glutamine/ 4% PFA/DEPC-PBS for 30’ at room temperature and then washed twice with DEPC-PBST. Samples were then incubated with 68°C pre-warmed Hyb for 15’ at room temperature after which Hyb was refreshed and structures incubated for 2 more hours at 68°C. If not immediately used for RNA probe hybridization, the samples were pre-cooled for 15’ at room temperature and stored at −20 °C. Prior to hybridization with the RNA probe, the samples and Hyb solution were pre-warmed at 68°C and incubated in fresh Hyb for 15’ at 68°C. Meanwhile, the RNA probe was diluted in Hyb (200ng/ml) and pre-heated for 13’ at 80°C in a heating block. The samples were then incubated with the RNA probe at 68°C overnight. The Hyb solution of the 15’ incubation step was stored at −20°C for the first washing step on the next day. All steps were performed under rocking.

#### Post-hybridization washes, antibody incubation and post-antibody washes

The next day, samples were washed once with Hyb (stored from day before) for 30’ at 68°C, twice with Solution 1 for 30’ at 68°C, twice with Solution 3T for 30’ at 68°C, twice with Solution 3T for 1h at 68°C and three times with TBST for 15’ at room temperature. During the 1h washes with Solution 3T, antibody solution was prepared. TBST with one grain of embryo powder was pre-heated for 30’ at 70°C in a water bath and cooled down on ice. Subsequently, 1% v/v lamb serum and 0.2% v/v Anti-DIG antibody (*Roche*) were added and incubated for 1h at 4 °C in the dark while rocking. The mix was centrifuged for 10’ at 4000rpm at 4°C and 1% lamb serum/TBST was added to the supernatant (final antibody concentration 1:2000). After finishing the washing steps, the samples were blocked with 10% lamb serum/TBST for 2.5h at room temperature and incubated with the antibody solution overnight at 4°C. All steps were performed under rocking. The next day, samples were washed twice with TBST for 15’, twice for 30’ and six times for 1h, all at room temperature. The final washing step was performed overnight in TBST at 4°C.

#### Staining

Samples were washed four times in freshly prepared NTMT for 15’ at room temperature under rocking. In the meantime, the staining solution BM Purple (*Roche*) was pre-warmed at room temperature and centrifuged for 1’ at 13.200 rpm. The supernatant was then used for staining the samples and incubated until a clear and specific signal appeared. The first 15’ of staining were performed under rocking, afterwards without. For stopping the staining reaction, the samples were washed once with NTT and twice with PBST, for at least 10’ each at room temperature under rocking. The samples were stored in 4% PFA/PBS at 4°C. Embryos, trunk-like-structures and gastruloids were imaged with the *AxioZoom.V16* (Zeiss).

### Single-cell transcriptome profiling of TLSs

96h, 108h TLS and 120h TLS were generated as described above. For 96h, 6 structures were selected based on the presence of a T^mCh+^ pole and absence of multiple axes formation; for 108h TLS, 5 structures were selected based on the presence of a T^mCh+^ pole, one axis of elongation, and initiation of neural tube formation, but no segmentation in the T^mCh+^ domain; for 120h TLS, 3 structures were selected based on the presence of a T^mCh+^ pole, one axis of elongation, clear formation of a neural tube Sox2^Ve+^ domain and segmentation in the T^mCh+^ domain. TLSs were picked with a p200 with the pipette tip cut-off at the 50µl mark, and serially washed through pipette transferring (cut 200µl tip) in wells filled with 200µl of 1xPBS/0.4%BSA (5 transfers) to get rid of the Matrigel. TLSs of the same time point were then pooled together and dissociated in 200µl TrypLE Express (Gibco) for 15 minutes (96h), 20 minutes (108h TLS) and 25 minutes (120h TLS) at 37°C, with pipetting every 5min intervals. The cell suspension was filtered using Scienceware Flowmi Cell Strainers, 40µm. Cells were washed twice with 1ml 1xPBS/0.4%BSA with centrifugation steps performed for 5 minutes at 1200rpm in low DNAbind Eppendorf tubes. The cell concentration was determined using a hemocytometer and cells were subjected to single-cell RNA sequencing (10x Genomic, Chromium™ Single Cell 3’ v3; one reaction per timepoint has been used) aiming for a target cell recovery of up to 10,000 sequenced cells per sequencing library (timepoint). Single-cell libraries were generated according to the manual, with one modification: fewer PCR cycles (*n*=8) were ran than recommended during cDNA amplification or library generation/sample indexing to increase library complexity. Libraries were sequenced with a minimum of 230 million paired end reads according to parameters described in the manual.

### Bulk RNA-seq of gastruloids and trunk-like-structures

96h aggregates, gastruloids, TLS, TLS^C^ and TLS^CL^ were generated as described above. We sequenced 3 biological replicates per condition (96h aggregates, Gastruloids, TLS, TLS^C^ and TLS^CL^). For 96h, 10 structures per replicate were selected (see previous section for selection criteria) and pooled; for Gastruloids, 6 structures per replicate have been selected (based on the presence of a T^mCh+^ pole and one axis of elongation) and pooled; for TLS, TLS^C^ and TLS^CL^, 6 structures per replicate were selected (see previous section for TLS selection criteria; TLS^C^ and TLS^CL^ were selected based on the presence of of a T^mCh+^ pole, elongation along one axis, segmentation in bunches of grapes at the anterior T^mCh+^ domain) and pooled. All samples were washed twice with 1xPBS/0.4%BSA. Then 350µl of RLT Plus buffer containing 1% β-mercaptoethanol (Thermo) was added to dissociate the structures and lyse the cells. After pipette dissociation and vortexing, samples were frozen at −80C. The next day, RNA was extracted using RNeasy Plus Micro Kit (Qiagen) and RNA concentration and quality was measured using the Agilent RNA 6000 Pico kit on an Agilent 2100 Bioanalyzer. All samples analyzed had a RINe value higher than 8.0, and were subsequently used for library preparation. mRNA libraries were prepared using KAPA Stranded RNA-Seq Kit (KapaBiosystem) according to the manufacturer’s instructions. 500ng of total RNA was used for each sample to enter the library preparation protocol. For adapter ligation dual indexes were used (NEXTFLEX® Unique Dual Index Barcodes NOVA-514150) at a working concentration of 71nM (5µl of 1uM stock in each 70µl ligation reaction). Quality and concentration of the obtained libraries was measured using Agilent High Sensitivity D5000 ScreenTape on an Agilent 4150 TapeStation. All libraries were sequenced using 75bp-paired end sequencing (150 cycles kit; FC-410-1002) on a HiSeq4000 platform at a minimum of 23,7 million fragments per sample.

### Expression profiling of TLS variability using NanoString

mESCs, mouse embryos and TLS were generated, selected and dissociated as described above. For mESCs, a total of 20,000 cells were harvested and frozen in 350µl RLT Plus buffer containing 1% β-mercaptoethanol (Thermo). Three embryos for each developmental stage (E8.5 and E9.5) were decapitated in order to obtain the post-occipital portion, pooled, dissociated, and frozen in 350µl RLT Plus buffer containing 1% β-mercaptoethanol. For 120h TLS, 9 individual structures have been selected (based on the selection criteria outlined before), dissociated independently and frozen in 350µl RLT Plus buffer containing 1% β-mercaptoethanol. RNA was isolated from the 12 samples in parallel using RNeasy Plus Micro Kit (Qiagen) and RNA concentration was measured using Qubit™ RNA HS Assay Kit.

To profile the expression of 41 genes and 4 housekeeping genes (*Polr1b*, *Hprt*, *Abcf1*, *Gusb*), a range of 36-161 ng total RNA/sample were used in a NanoString nCounter Element assay to profile the 12 individual samples. Probe hybridization was set up according to manufacturer’s instructions and performed for 24 hours (MAN-10040-05). Reactions were ran on the NanoString nCounter SPRINT Instrument. False negative probes detected up to 5 counts, which informed the magnitude of potential false negative signal. Thus, 6 counts were conservatively removed from all measurements. Positive control probes normalization step was applied (geometric mean) and finally the combination of the four housekeeping genes was used to obtain the final normalized counts table. To assess variability in the expression of individual marker genes, the log_2_(MAX(all_samples)/MIN(all_samples)) was used as a proxy with a high value representing high variability and vice versa. See **Supplementary Table 6** for probes design and sequences and Data S7 for normalized gene counts.

### Computational analysis

If not stated otherwise: All statistics and plots are generated using R version 3.6.0 “Planting of a Tree” and Seurat version 3.0^54^.

### Single-cell transcriptome profiling of TLSs

#### Preprocessing

The Cell Ranger pipeline version 3 (10x Genomics Inc.) was used for each scRNA-seq data set to de-multiplex the raw base call files, generate the fastq files, perform the alignment against the mouse reference genome mm10, filter the alignment and count barcodes and UMIs. Outputs from multiple sequencing runs were also combined using Cell Ranger functions.

#### Quality control

The initial quality control was performed using scanpy^55^. Cells with less than 10,000 or more than 40,000 counts, a mitochondrial-fraction above 0.1 and less than 3,000 genes were flagged as insufficient.

#### Cluster determination

Single cell data created for the three developmental time points (96h, 108h and 120h) were loaded to Seurat^54^, with a minimum requirement of 200 features and 3 cells and filtered for previously flagged barcodes. Subsequently, the expression data were independently normalized, variable features were detected and log-normalized and scaled to 10,000 (default settings). Next, for downstream integration of the three time points sequenced, a PCA was run for each time point prior to integration anchor set detection (reduction = “rpca”, dims = 1:30). Finally, these integration anchors were used to integrate the data points using the previously calculated anchor sets. A list of cell cycle markers loaded with Seurat was used to cell cycle score all cells and subsequently run the default workflow for scaling with vars.to.regress set to cell cycle scores for S and G2M phase (**Extended Data** Fig. 15**a**). For downstream analysis and visualization of the integrated dataset, a PCA was run to then calculate a joint UMAP (dims = 1:30, n.neighbors = 10). Standard workflow steps were applied for cluster generation (FindNeighbors, dims = 1:20 and FindClusters, resolution = 0.5), resulting in a total of 15 clusters. Finally, two small clusters (Seurat_10 and Seurat_11) were removed due to presence of stressed cells (high mitochondrial RNA counts and low total RNA counts) (Seurat_10; **Extended Data** Fig. 15**b**) and doublets (shown by almost double amount of total RNAs counts and UMIs) (Seurat_11; **Extended Data** Fig. 15**c**). All remaining clusters show a similar distribution of average UMIs and genes detected per cell (**Extended Data** Fig. 15**d**). See **Supplementary Table 8** for marker genes list for each identified cluster.

#### Subclustering of somitic and neural cells

For subclustering of somitic cells, data were first split by sampled timepoints and somitic clusters (“Somite (0)”, “Somite”, “Somite (Dermo)”, “Somite (Sclero)”) were extracted. Subclustering was then performed in Scanpy (resolution=0.3). For subclustering of the neural tube cluster, cells assigned to “NeuralTube2” from TLS were extracted. Subclustering was then performed in Scanpy (resolution=0.65).

#### Comparison with mouse embryo

A previously established reference data set of mouse post-implantation development (E6.5 to E8.5)^35^ was utilized for gene expression comparison to *in vivo* data as well as cell type and proportion comparisons. The *in vivo* data were filtered to include only the relevant embryo time points (E.7.5, E8.0 and E8.5) all extra-embryonic and occipital cell-types (that are not generated in gastruloids^3^ and TLS (this study)) were removed. See **Supplementary Table 8** for marker genes list for each identified cluster. Both, the mouse *in vivo* and the TLSs *in vitro*, data sets were normalized in parallel (SCTransform, default settings). To adjust for different resolutions in cell type detection between the embryo and TLS datasets, the *in vivo* “somites” and “PSM” cell states were subclustered (resolution=0.15) to identify a cell cluster with an anterior PSM (aPSM) signature (**Extended Data** Fig. 15**e,f**). These cells were relabeled accordingly and subsequently treated as an individual cell type. The subclustering step also generated two clusters from the *in vivo* PSM original cell state, but due to marker genes similarity this subclustering was not taken into account in the further downstream analysis (**Extended Data** Fig. 15**e,f**). Prediction scores were used as measurement for certainty in cell type matching (“certainty scores”). Scores shown are means across all cells even if not assigned to the respective cluster. Finally, the *in vivo* mouse and *in vitro* TLS cell types were matched using an integrated reference, by first finding of transfer anchors, followed by data transfer using the anchors and finally the predicted cell types were used as *in vivo*/*in vitro* matched cell type counterparts. For differential expression and conserved marker calculation both data sets were merged and genes located on the sex chromosomes were excluded to avoid biases resulting from comparison of only male TLSs with *in vivo* cells from male and female embryos.

#### Analysis of conserved marker genes

The conserved marker genes was calculated on the integration of the TLS and embryo data as described above using the FindConservedMarkers function with default setting. Cut-offs used were set to a minimum percentage difference of 25% (for percent of cells expressing the gene of interest (GOI)), and an absolute minimum log_2_FC of 1. See **Supplementary Table 9** for list of top 25 conserved marker genes.

#### Analysis of differentially expressed genes between TLS and embryo

TLS and embryo data were integrated as described above. For the analysis, the *in vivo* cell states were used as a reference, and compared to the TLS cells attributed to that cluster. To this end, the percent of cells expressing a gene as well as the average expression for that gene in the *in vivo* cell states as well as the *in vitro* cells attributed to that cluster was computed using the FindMarkers function with default settings. Differential expression between TLS and embryo was tested for either all the genes expressed in the cluster of interest, or the 20 top marker genes of each *in vivo* cell states. Cut-offs used were a minimum percentage difference of 25% (for percent of cells expressing the gene of interest (GOI)), and an absolute minimum log_2_FC of 1. See **Supplementary Table 9** for list of genome wide differentially expressed marker genes.

#### RNA velocity and pseudo-time analysis

RNA velocity was calculated using the velocyto tool^31, 56^ and visualized using scanpy^55^. Trajectory inference and pseudo-time analysis were done using the scanpy package PAGA^57^. The previously calculated UMAP and cell cluster assignment was used for velocity projection and trajectories for pseudo-time. For pseudo-time analysis, somitic clusters (“Somite (0)”, “Somite”, “Somite (Dermo)”, “Somite (Sclero)”) were merged, the putative NMP cluster was set as root and 108h and 120h cells were analyzed separately.

## Bulk RNA-seq of gastruloids and trunk-like-structures

### Processing and expression levels

RNAseq triplicates of the different growth conditions 96h, Gastruloids 120h, TLS, TLS^C^ and TLS^CL^ were pre-processed using cutadapt^58^ to remove adapter and trim low quality bases. Reads were subsequently aligned against the reference genome mm10 using STAR^59^ (parameter: outSAMtype BAM SortedByCoordinate --outSAMattributes Standard --outSAMstrandField intronMotif --outSAMunmapped Within --quantMode GeneCounts). Finally, Stringtie^60^ was used for transcript assembly, e.g. calculation of strand-specific TPMs. The three biological replicates highly correlate between each other (**Extended Data** Fig. 15**g**).

### Differential gene expression

Differential expression analysis was done independently per group comparison using the R package DESeq2^61^ using the raw expression counts from STAR’s reads per gene output. *z*-scores were calculated according to the formula ((VALUE(sample)-AVERAGE(all_samples))/STDEV(all_samples). TPM values for all genes can be found in **Supplementary Table 10**.

### Gene Set Enrichment Analysis (GSEA)

GSEA was performed using the R package “fgsea”^62^ (minSize=5, maxSize=500, number of permutations = 10.000) taking all genes statistically differentially expressed between TLS and gastruloids (FDR < 0.05, no log_2_FC cut-off).

### Data availability

Bulk and single-cell RNA-Seq data have been deposited in the Gene Expression Omnibus (GEO) under accession code GSE141175.

### Code availability

All computational code used in this study is available upon request.

---

**Supplementary Video 1.**

Wide-field live imaging of TLS from 96-120h showing elongation, somite segmentation, and neural tube formation.

**Supplementary Table 1.**

Lists of genes used to generate boxplots from Figure 2b and **Extended Data Fig. 3e**

**Supplementary Table 2.**

Significantly enriched processes in TLS versus Gastruloids as revealed by Gene Set Enrichment Analysis

**Supplementary Table 3.**

Overview of buffers, culture media, kits, and antibodies used in this study

**Supplementary Table 4.**

Sequences of oligo’s and *in situ hybridization* probes used in this study

**Supplementary Table 5.**

Raw Ct values from quantitative PCR analysis

**Supplementary Table 6.**

Details of Nanostring probes used in this study

**Supplementary Table 7.**

Nanostring normalized gene counts

**Supplementary Table 8.**

Marker genes for TLS clusters (single-cell RNA-Seq) & Marker genes for embryo clusters (single-cell RNA-Seq)

**Supplementary Table 9.**

Top 25 conserved cluster marker genes between TLS and embryo & Genes differentially expressed between TLS cell types and their corresponding *in vivo* cell type

**Supplementary Table 10.**

Expression values in TPM (Transcripts Per Million) for all genes (bulk RNA-Seq)

**Extended Data Fig.1.**
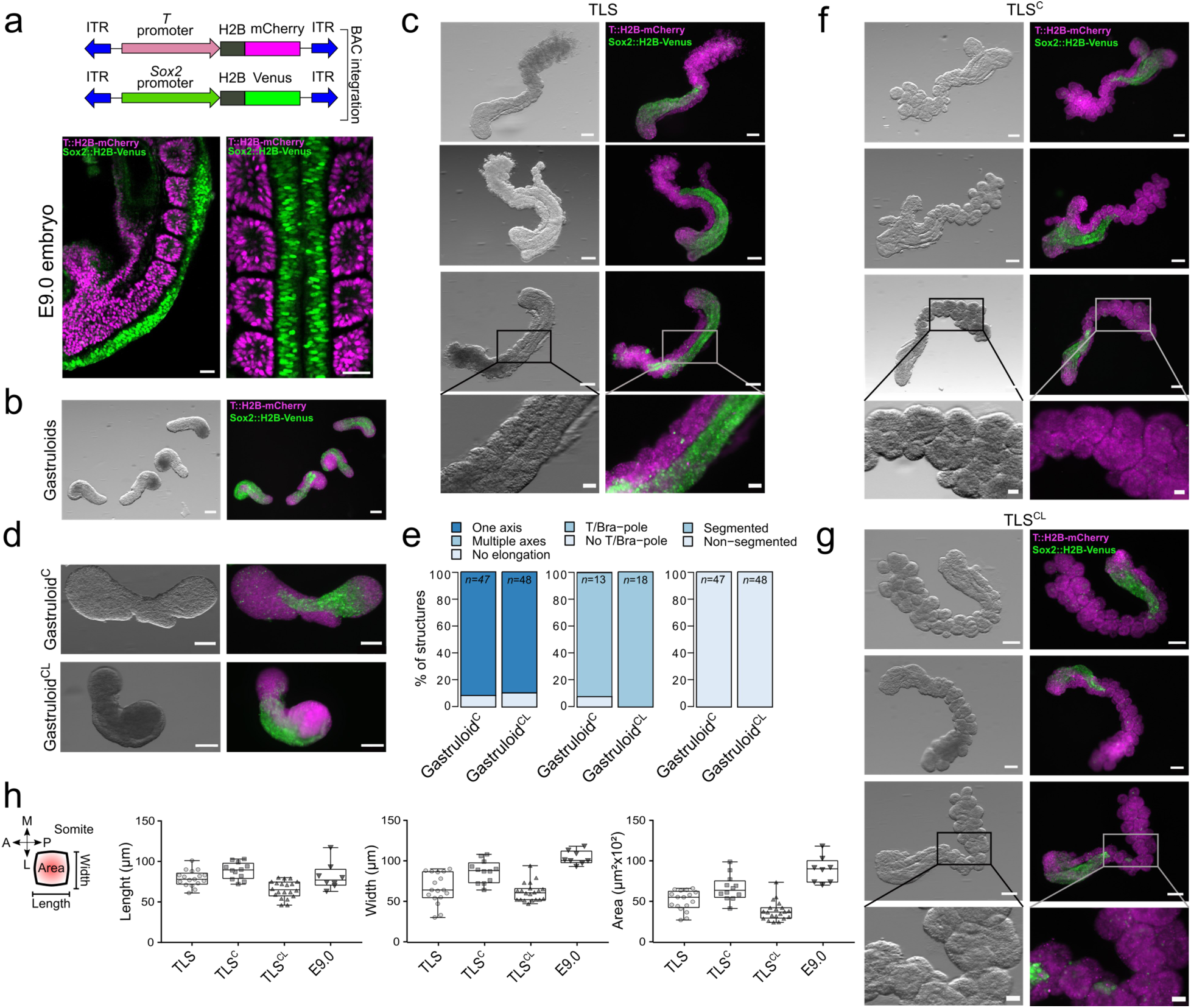
Induction of embryo-like morphology in trunk-like-structures. **a,** Schematic of *T::H2B-mCherry*; *Sox2::H2B-Venus* double reporter mESCs (upper panel) and *in vivo* validation by tetraploid complementation (bottom panel). Optical sections of light-sheet imaging of a E9.0 mouse embryo caudal end (bottom-left panel) and of somites and neural tube (bottom-right panel). Scale bar 50μm. **b**, Wide-field fluorescent and bright-field images of 120h gastruloids. Scale bar 200μm. **c**, Wide-field fluorescent and bright-field images of 120h TLS. **d**, Wide-field fluorescent and bright-field images of 120h gastruloids treated with CHIR or CHIR+LDN from 96 to 120h showing no signs of segmentation in the absence of 5% matrigel. **e**, Quantifications of gastruloids from conditions in **d**. Treatment of gastruloids with CHIR and LDN does not significantly alter the axis formation and induction of a T^mCH+^ pole, and is not able to induce segmentation. **f**, Wide-field fluorescent and bright-field images of 120h TLS^C^. **g**, Wide-field fluorescent and bright-field images of 120h TLS^CL^. In **b-d** and **f,g**, scale bar is 200μm, except for magnifications, where scale bar is 50μm. **h**, Quantification of somite size in TLS and embryos using Fiji (see **Supplemental Information**) reveals that TLS and TLS^C^ somites have a similar length but a slightly smaller width as compared to the embryo, resulting in an overall slightly reduced segment area. In TLS^CL^, both somite length and width are smaller than *in vivo*. Boxes indicate interquartile range. End of whiskers represent minimum and maximum. Symbols indicate individual somite. Central line represent the median.

**Extended Data Fig.2.**
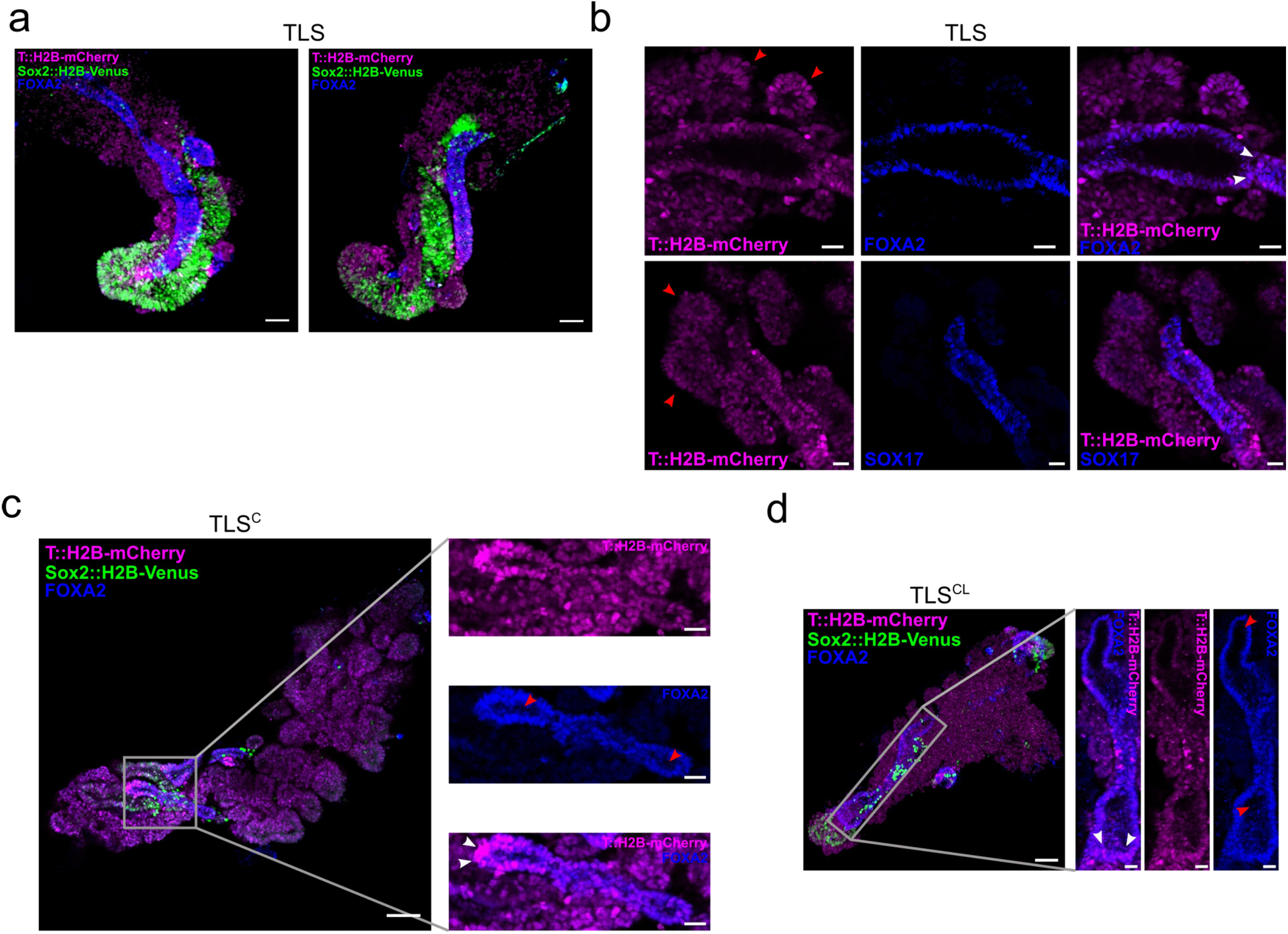
Formation of gut-like-structures in trunk-like-structures. **a,** 3D maximum intensity projection of TLS (left panel) stained for FOXA2 (blue) to reveal the gut-like-structure. Scale bar 50μm. **b,** Confocal sections showing the FOXA2^+^ (upper panel) and SOX17^+^ (bottom panel) gut-like structure in TLS. Note FOXA2^+^/T^mCH-high^ cells at the base of the gut (white arrowheads). Red arrowheads indicate somites. Scale bars 25μm. **c,** Confocal section showing gut-like structure in TLS^C^. **d,** 3D maximum intensity projection showing gut-like structure in TLS^CL^. In the magnifications (right panels) in **c,d** white arrowheads indicate FOXA2^+^/T^mCH-high^ cells at the base of the gut. Red arrowheads indicate gut tubular cavity.

**Extended Data Fig.3.**
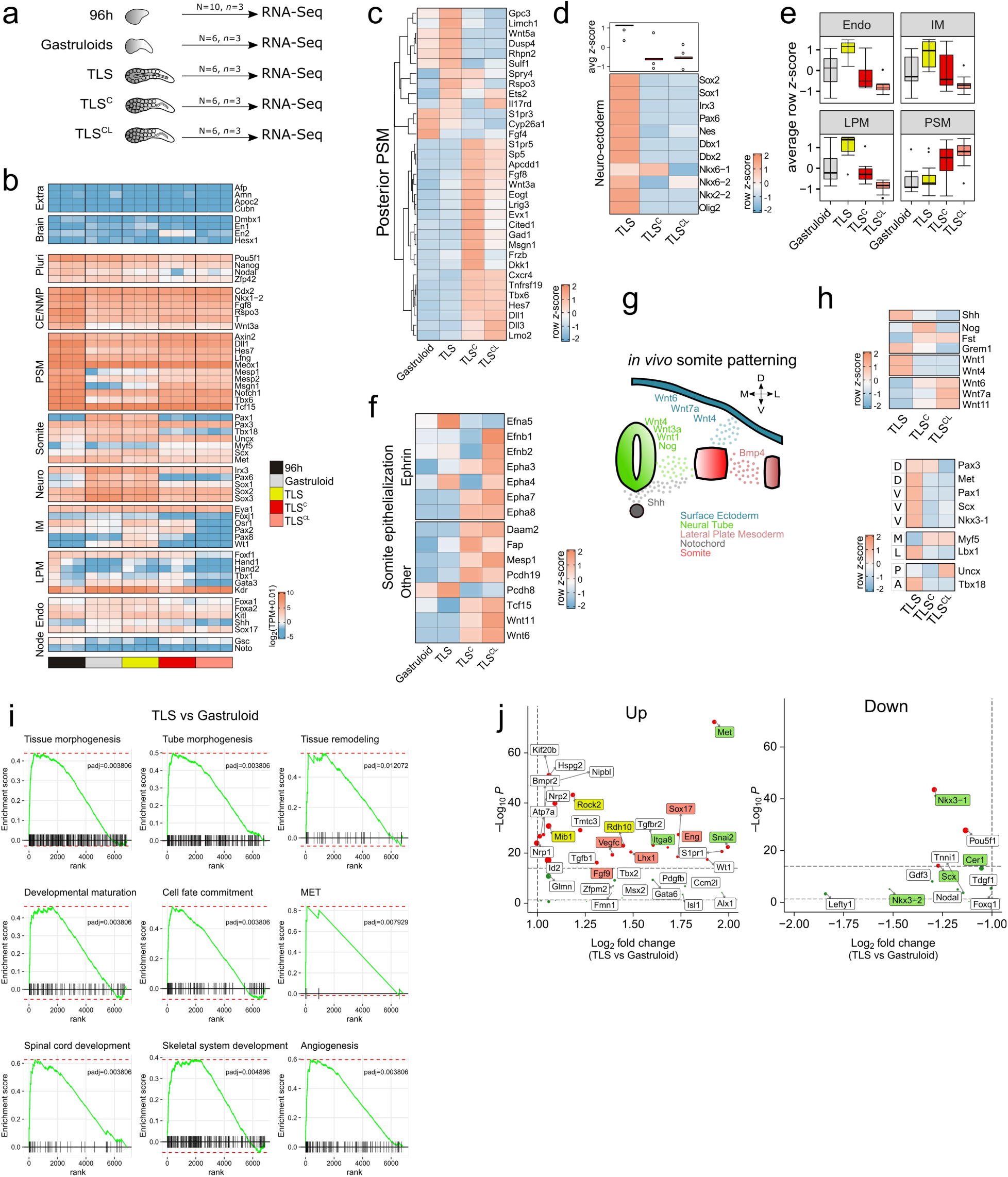
Bulk RNA-Sequencing of gastruloids and trunk-like-structures. **a,** Schematic of experimental set-up. N=number of pooled structures per replicate, *n*=number of replicates. **b,** Heatmap of expression for an extended panel of selected genes associated with development of indicated embryonic structures in 96h aggregates, gastruloids, TLS, TLS^C^ and TLS^CL^, as measured by RNA-seq. TPM, Transcripts Per Million. CE, caudal end, NMP, neuromesodermal progenitors, PSM, presomitic mesoderm, LPM, lateral plate mesoderm, IM, intermediate mesoderm. **c,** Heatmap with scaled expression (row *z*-score) of known marker genes of posterior presomitic mesoderm (pPSM)^8^. Scores are an average of the 3 independent replicates per condition. **d**, Heatmap with scaled expression (row *z*-score) of neural tube known marker genes. Scores are an average of the 3 independent replicates per condition. Boxplot shows distribution of *z*-scores (per column) for different conditions. Dots indicate outliers. **e,** Boxplots showing distribution of *z*-scores for known marker genes of indicated tissue layers. Endo, Endoderm; IM, Intermediate Mesoderm; LPM, Lateral Plate Mesoderm; PSM, Pre-Somitic Mesoderm. Boxes indicate interquartile range. End of whiskers represent minimum and maximum. Dots indicate outliers. Central line represent the median. List of genes used for each category in **Supplementary Table 1. f,** Heatmap of scaled expression (row *z*-score) of somite epithelialization factors in gastruloids, TLS, TLS^C^ and TLS^CL^. Scores are an average of the 3 independent replicates per condition. **g,** Schematic overview of signalling factors involved in somite compartmentalization *in vivo*. M, medial; L, lateral; D, dorsal; V, ventral. **h,** Heatmap with scaled expression (row *z*-score) of signalling factors depicted in **g** (top panel) and heatmap with scaled expression of marker genes for different somite compartments (bottom panel). A, anterior; P, posterior; M, medial; L, lateral; D, dorsal; V, ventral. Scores are an average of the 3 independent replicates per condition. **i,** Gene Set Enrichment Analysis enrichment plots of selected significant pathways. **j,** Up- and downregulated genes involved in embryo & tissue morphogenesis (related to Fig. 2e). Dot size scales with log_2_ of absolute expression. Red dots, log_2_FC>2 and padj<10e^-15^. Green dots, log_2_FC > 2. Upper dotted line, padj=10e^-15^; bottom dotted line, padj(FDR)=0.05. Green label, involved in somitogenesis; orange label, involved in blood vessel development; yellow label, involved in both.

**Extended Data Fig.4.**
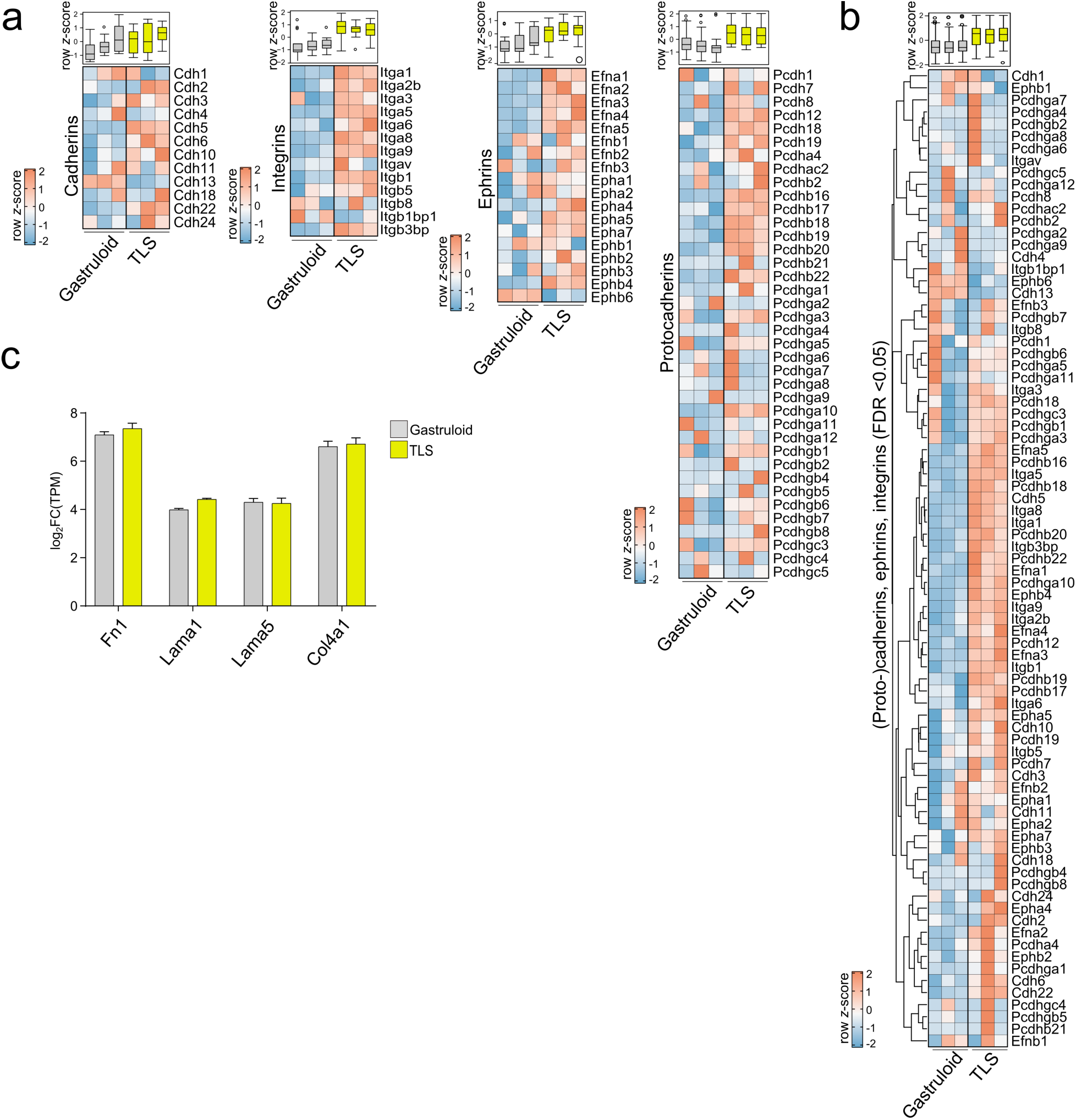
Expression of cell adhesion molecules in gastruloids and TLS. **a**, Heatmap with scaled expression (row *z*-score) of cadherins, protocadherins, and ephrins in gastruloids and TLS. Boxplots show distribution of *z*-scores (per column) for different samples. Boxes indicate interquartile range. End of whiskers represent minimum and maximum. Dots indicate outliers. Central line represent the median. Every column in each heatmap represent one of the three biological replicates. **b,** Heatmap of scaled expression of genes in **a** with significantly differential expression (FDR<0.05). **c,** Bar graph showing expression of integrin ligands fibronectin (Fn1), laminin alpha-1 (Lama1), laminin alpha-5 (Lama5) and collagen 4a1 (Col4a1) in gastruloids and TLS. Bar graphs represent average of 3 samples. Error bars represent standard deviation.

**Extended Data Fig.5.**
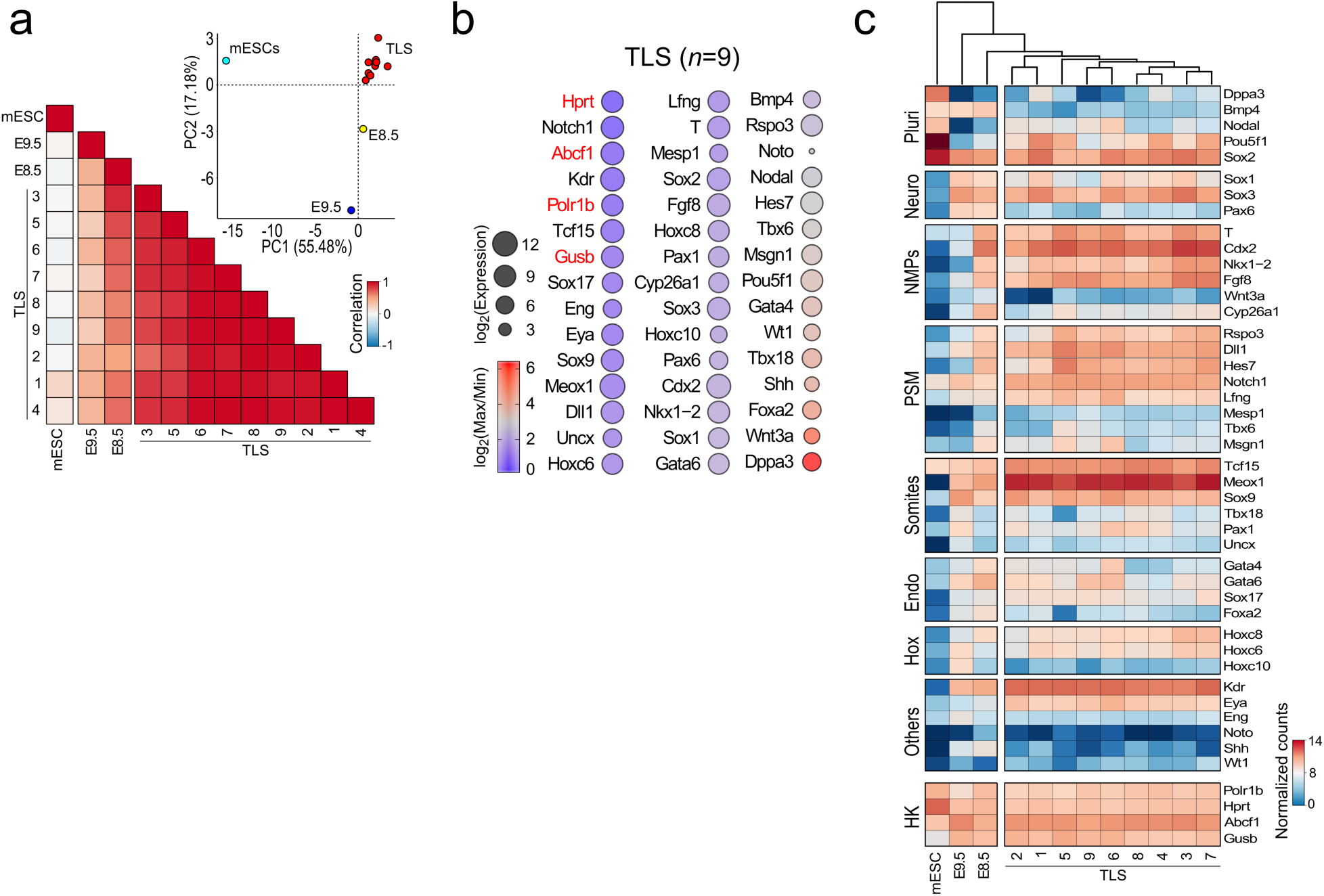
NanoString analysis of individual TLS. **a,** Heatmap correlation plot and PCA analysis of mESCs, E8.5 and E9.5 post-occipital embryo, and 9 individual TLS based on expression of 41 developmental genes measured by NanoString analysis. PC1 and PC2 represents the two components with highest percentage of explained variance. **b,** Dot plot for 45 genes (41 test genes and 4 housekeeping genes (red font)). Dot size scales with log_2_ of expression. Color indicates log_2_(max/min) as a proxy for range of expression. Genes are ranked from lowest range (upper left) to highest range (bottom right). **c,** Heatmap with normalized counts of indicated *in vivo* marker genes for embryonic cell types in mouse embryonic stem cells (mESCs), the E8.5 and E9.5 post-occipital embryo, and 9 individual TLS. Pluri, pluripotency; Neuro, neuro-ectoderm; NMPs, neuromesodermal progenitors; PSM, presomitic mesoderm; Endo, endoderm; Hox, Hox genes; HK, housekeeping genes.

**Extended Data Fig.6.**
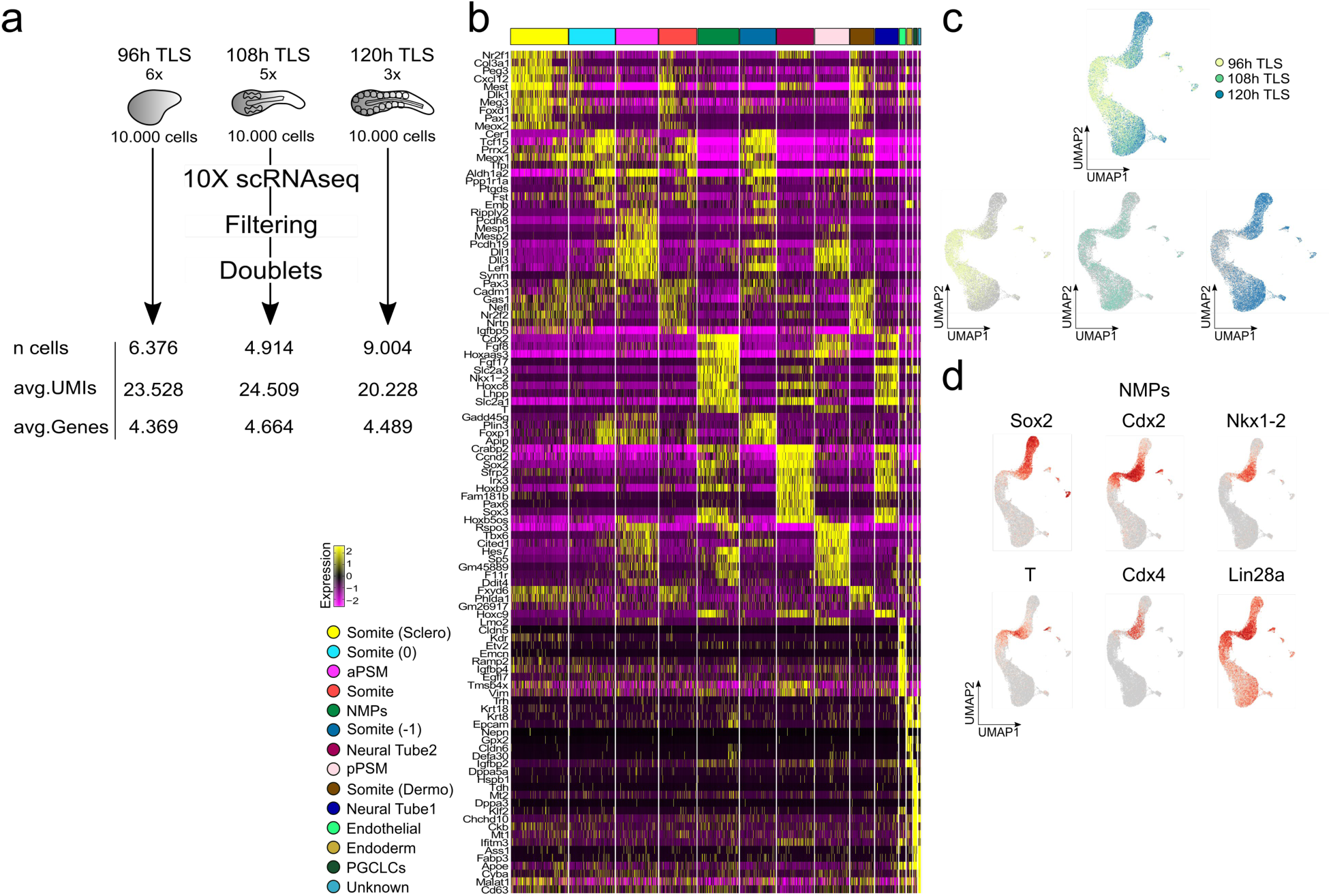
Single-Cell RNA-sequencing of TLS. **a,** Experimental set-up. Criteria for filtering and removal of doublets are described in **Methods**. **b,** Heatmap with scaled expression of top marker genes of the fourteen identified clusters. **c,** UMAPs coloured by sampled timepoints. **d,** UMAPs coloured by expression of indicated *in vivo* marker genes for neuromesodermal progenitors (NMPs). pPSM, posterior pre-somitic mesoderm; aPSM, anterior pre-somitic mesoderm; PGCLCs, primordial germ cell like cells.

**Extended Data Fig.7.**
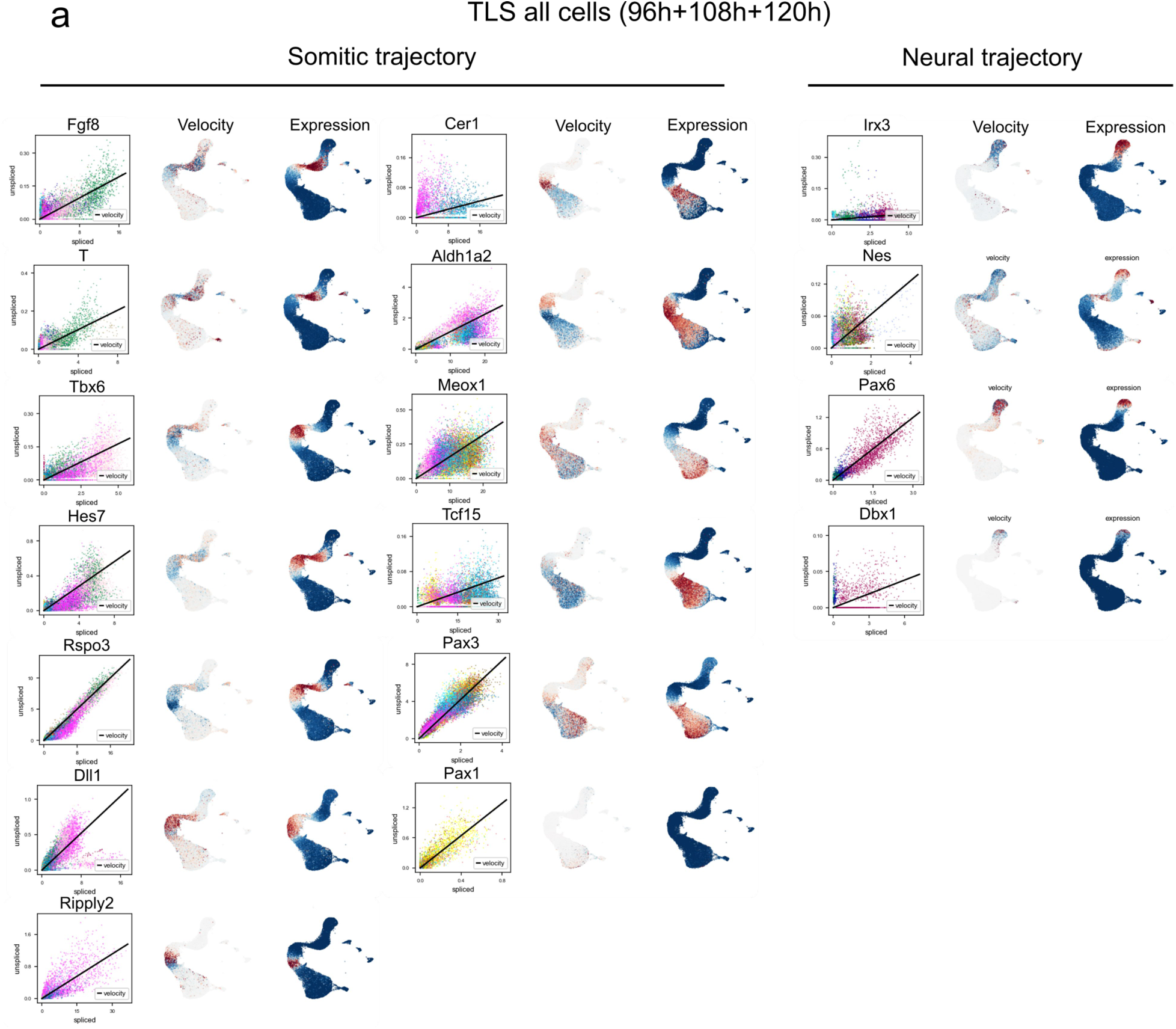
RNA Velocity of *in vivo* cell type marker genes in TLS. **a,** Velocity and expression of *in vivo* marker genes of the somitic and neural trajectory projected on the pooled (96+108+120h TLS) UMAP. Cells are coloured following the cluster colours in Fig. 3a.

**Extended Data Fig.8.**
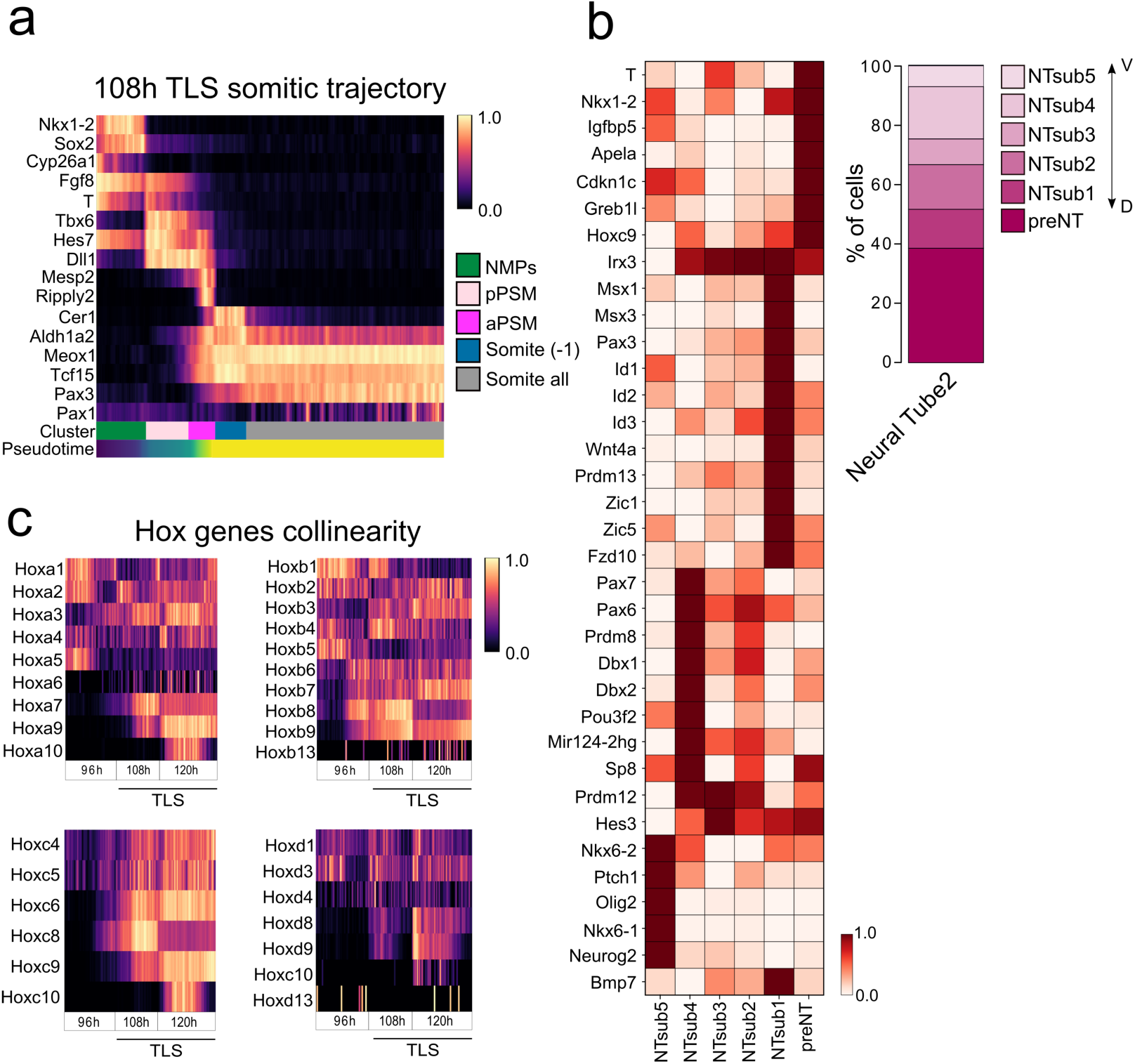
Spatiotemporal progression of gene expression in TLS. **a**, Heatmap with scaled expression of genes involved in somitogenesis in 4914 cells from 108h TLS rooted in NMPs and ordered by pseudotime. aPSM, anterior PSM, pPSM, posterior PSM. **b,** Heatmap with scaled expression for neural lineage marker genes of each cluster that confer progenitor identity along the DV-axis *in vivo* (left panel) and column plot showing subcluster distributions. D, Dorsal; V, Ventral. **c,** Heatmap with scaled expression of Hox genes in NMPs and their direct descendants (Neural Tube1, pPSM), ordered by pseudotime after rooting in 96h.

**Extended Data Fig.9.**
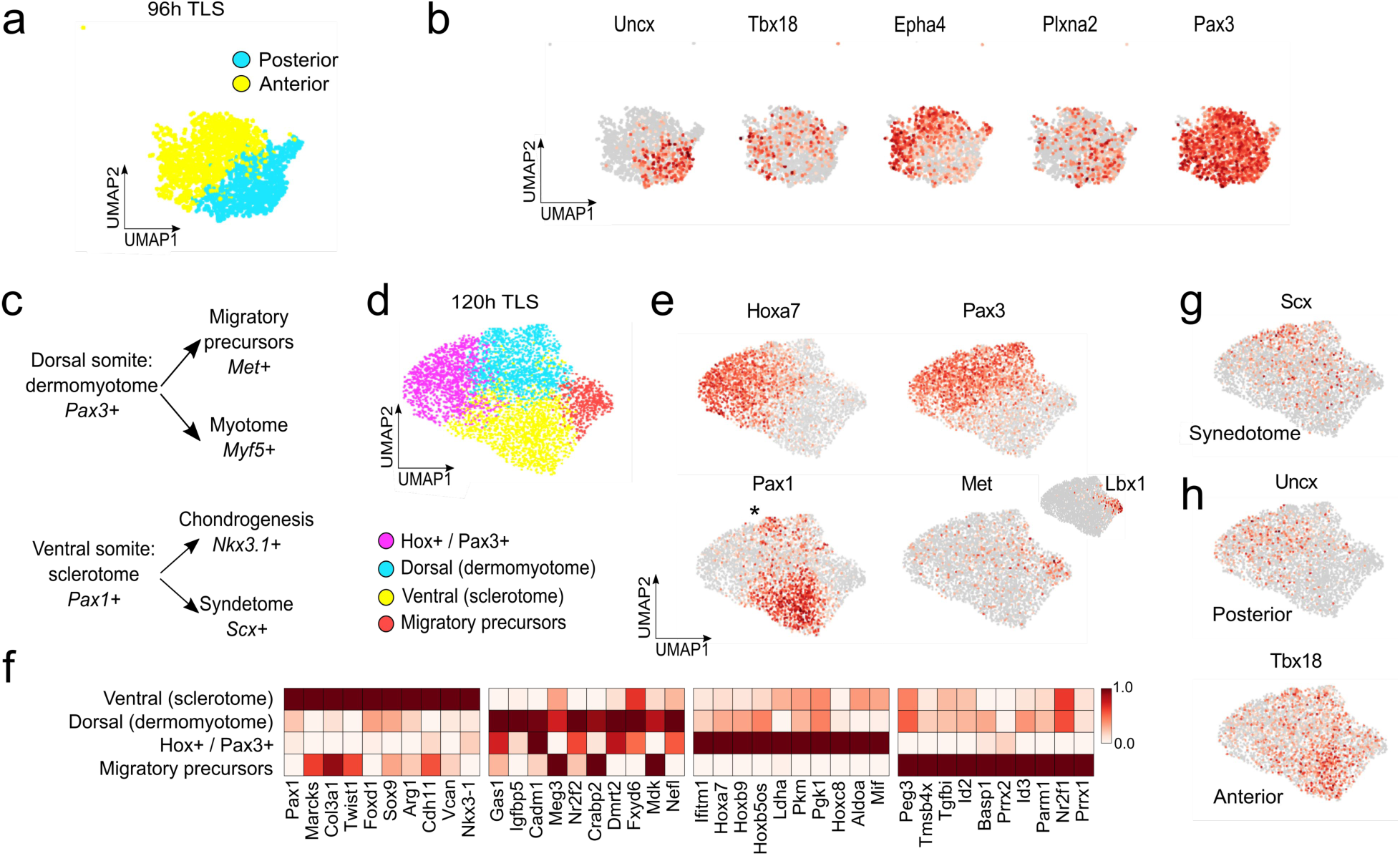
Somite compartmentalization in TLS. **a,** UMAP representation of reclustering of 96h TLS somitic cells. **b,** UMAP coloured by expression of indicated *in vivo* marker genes for anterior and posterior somite compartments. **c,** *In vivo*, the dorsal somite compartment is marked by Pax3 expression and gives rise to dermamyotome and subsequently migratory muscle precursors and myotome. The ventral somite compartment is marked by Pax1 and gives rise to the sclerotome, which subsequently differentiates into chondrocytes and syndetome. **d,** UMAP representation of reclustering of 120h TLS somitic cells. **e,** UMAP coloured by expression of indicated marker genes for dorsal and ventral somite compartments. Asterisk refers to Pax1^+^/Scx^+^ population. **f,** Heatmap with scaled expression for top 10 marker genes of each cluster. **g,** UMAP coloured by expression of *in vivo* syndetome marker Scx. Note that the Scx domain partially overlaps with a separate cluster of Pax1^+^ cells (indicated with an asterisk in **e**). **h,** UMAP coloured by expression of posterior (upper panel) and anterior (bottom panel) markers Uncx and Tbx18.

**Extended Data Fig.10.**
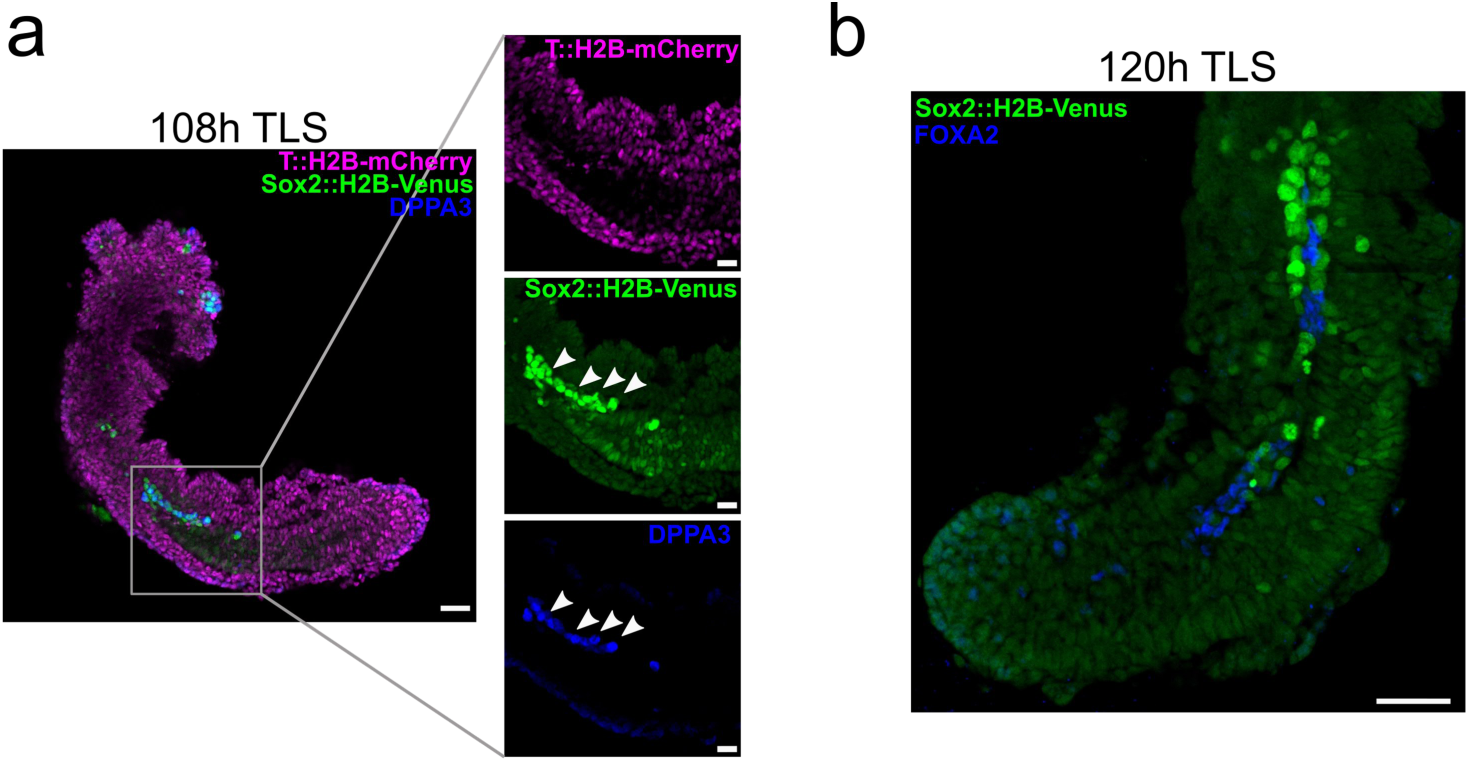
Primordial Germ Cell Like Cells (PGCLCs) in TLS. **a,** Confocal section of 108h TLS showing PGCLCs that co-express Sox2^VE^ and DPPA3. **b,** Confocal section of 120h TLS stained for FOXA2 and Sox2^VE^. Note how Sox2^VE-high^ cells contact the FOXA2^+^ cells in 120h TLS. Scale bars 50μm, 25μm for magnifications.

**Extended Data Fig.11.**
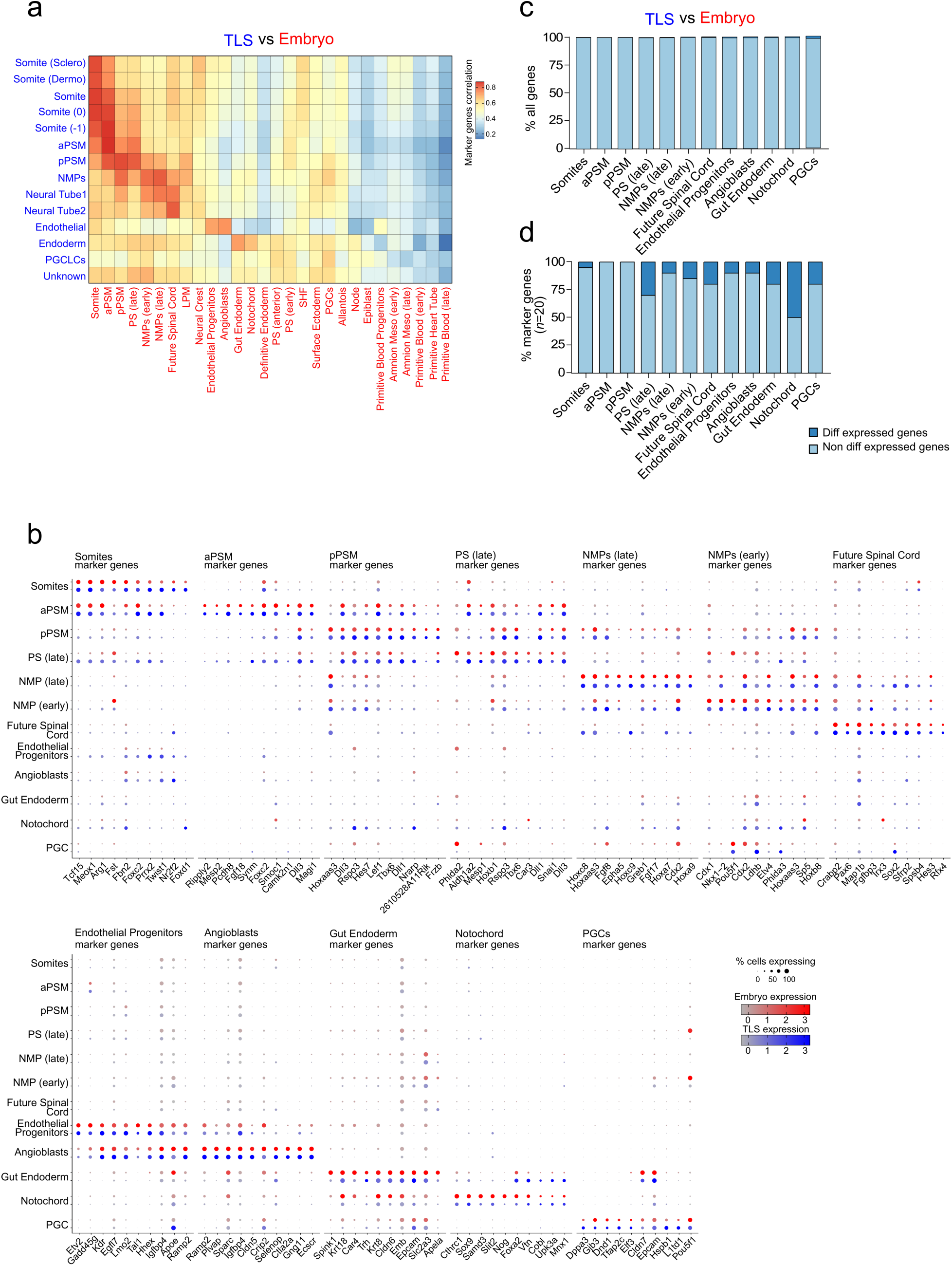
Comparison of TLS cell-types with post-occipital cell-types of the mouse embryo. **a,** Heatmap showing correlation of cell types identified in TLS (blue font) and cell types identified in the post-occipital E7.5 and E8.5 embryo (red font) base on top 20 marker genes for *in vivo* clusters. **b,** Dot plots showing the ten most conserved cell type markers for indicated cell types between TLS and the embryo. **c,** Bar graph with percentage of expressed genes (average expression in cluster > 0) that are differentially expressed in TLS as compared to the embryo (min.diff.pct > 0.25 and log_2_FC > 1). **d,** Bar graph with percentage of top 20 *in vivo* cluster marker genes that are differentially expressed in TLS as compared to the embryo (min.diff.pct > 0.25 and log_2_FC > 1).

**Extended Data Fig.12.**
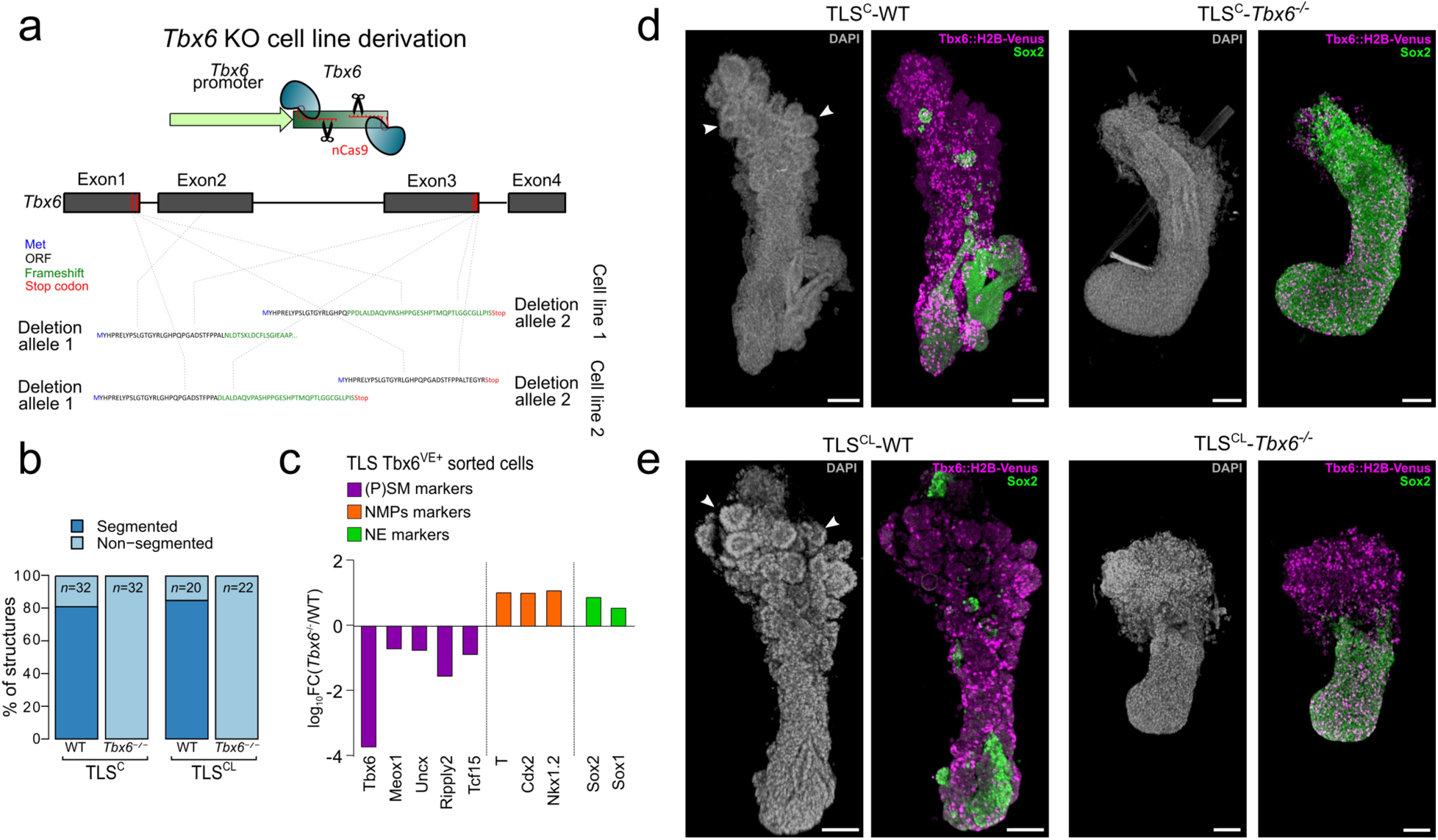
Loss of segmentation and molecular phenotype in Tbx6^-/-^ TLS models. **a,** Schematic and Sanger sequencing validation of two independent *Tbx6::H2B-Venus* (Tbx6^VE^); *Tbx6^-/-^* mESC lines. **b,** Segmentation phenotype of TLS^C^-WT, TLS^C^-*Tbx6^-/-^*,TLS^CL^-WT and TLS^CL^*-Tbx6^-/-^*. Data represent 3 different experiments performed with 2 independent mESC lines of each genotype. **c,** qRT-PCR showing expression of (P)SM, NMPs and NE genes in sorted Tbx6^VE+^ cells from TLS-WT and TLS-Tbx6^-/-^. Log_10_FC was calculated from the fold change ratio between TLS-Tbx6^-/-^ and TLS-WT. Bars represent the mean of two biological replicates. **d,** 3D maximum intensity projections of TLS^C^-WT and TLS^C^-*Tbx6^-/-^* immunostained for Tbx6^VE^ (magenta) and SOX2 (green). **e,** 3D maximum intensity projections of TLS^CL^-WT and TLS^CL^*-Tbx6^-/-^* immunostained for Tbx6^VE^ (magenta) and SOX2 (green). Scale bars 100μm.

**Extended Data Fig.13.**
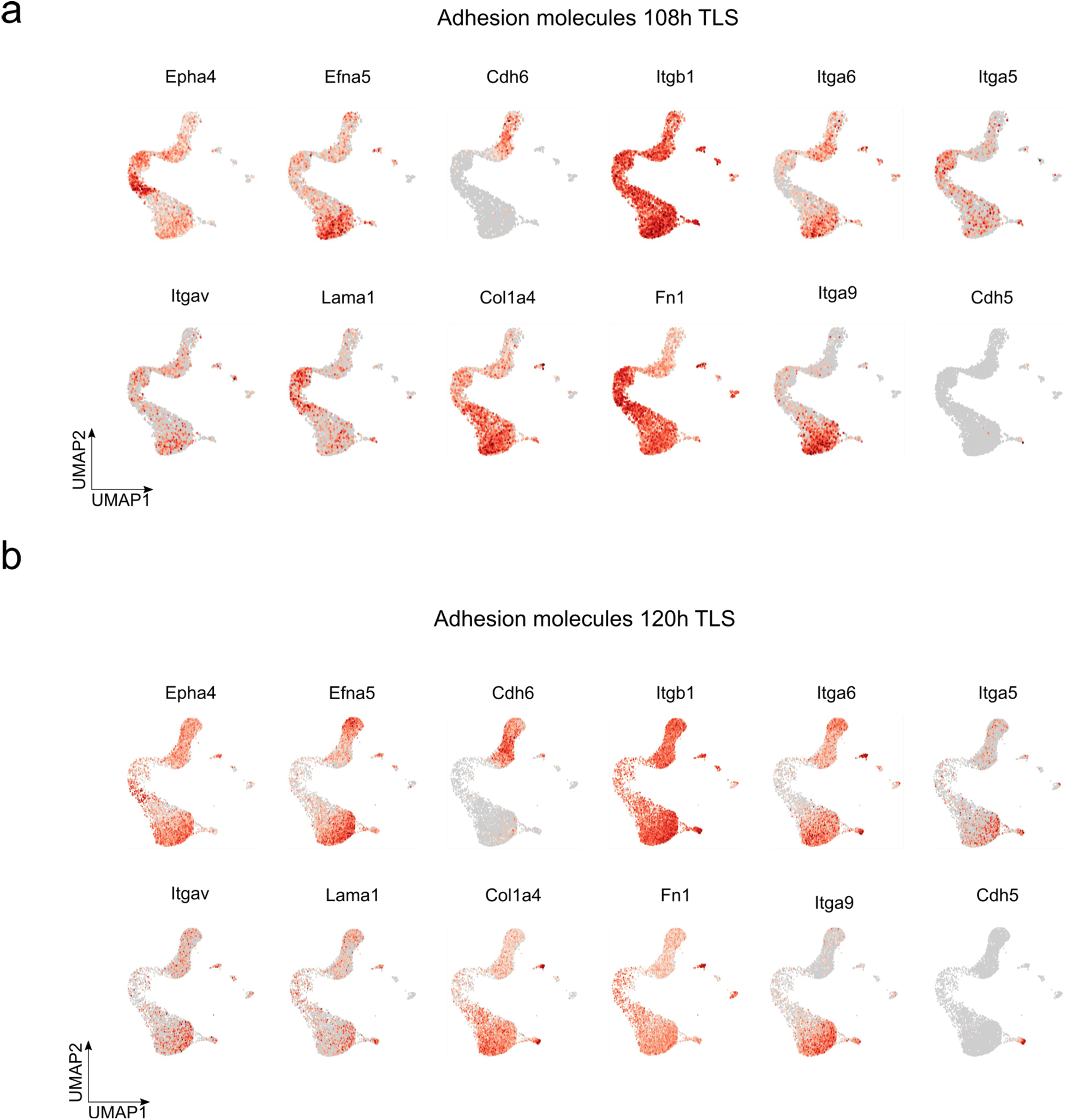
Cell type specific expression of cell adhesion molecules in TLS. **a,** UMAP of 108h TLS coloured by expression of indicated genes. **b,** UMAP of 120h TLS coloured by expression of indicated genes.

**Extended Data Fig.14.**
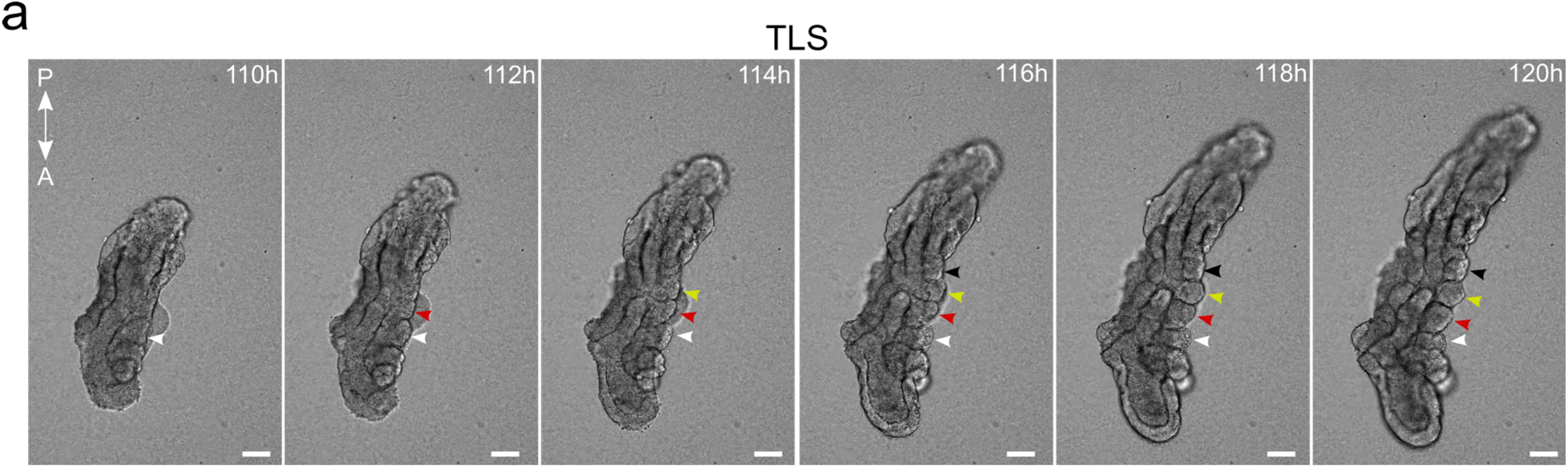
Sequential segmentation in TLS somites. **a,** Stills from brightfield live imaging showing sequential generation of somites along the anterior-posterior axis with approximately a two hour interval in TLS. Arrowheads indicate individual somites throughout time. Scale bar 100μm. A, anterior; P, posterior.

**Extended Data Fig.15.**
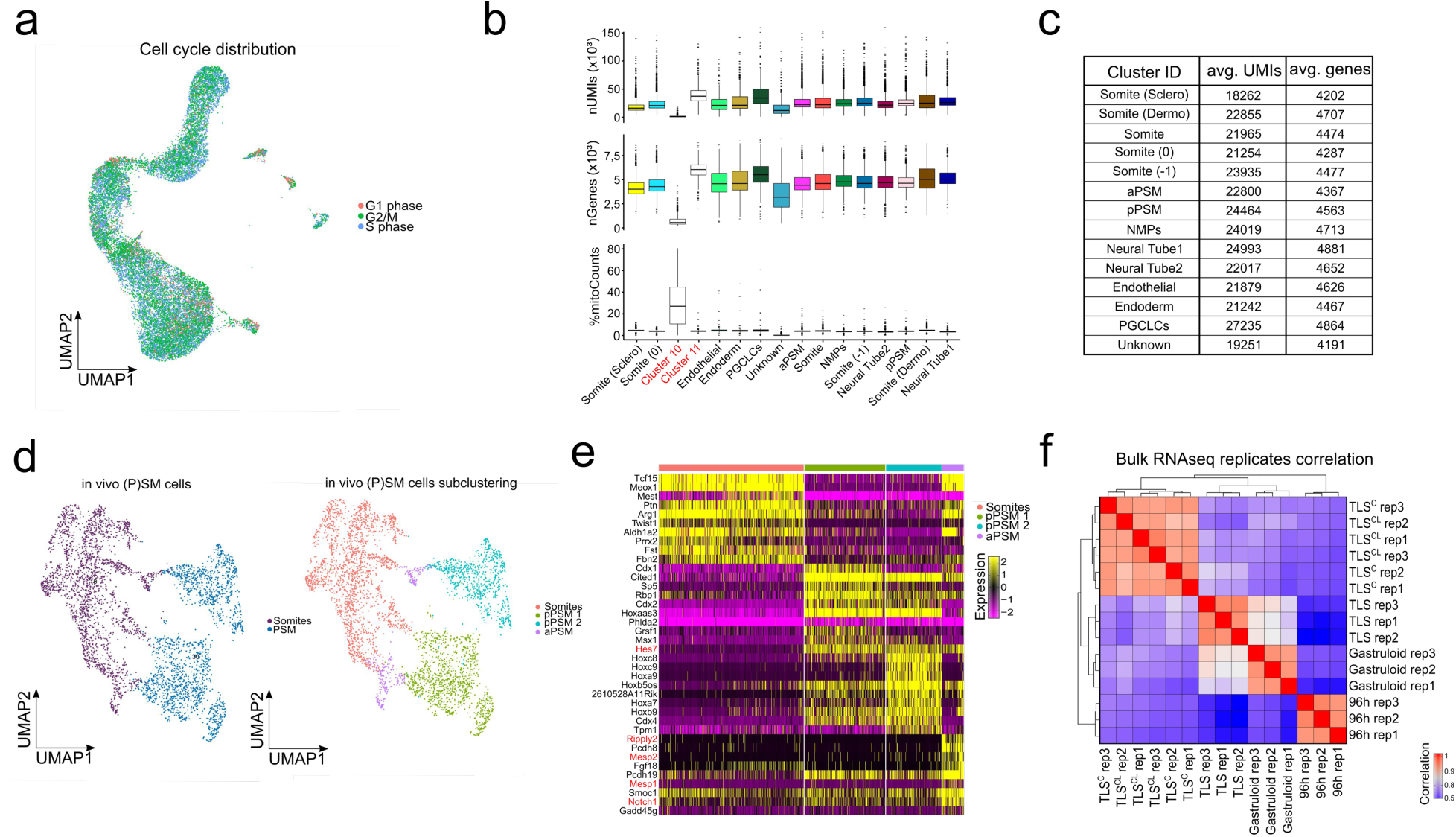
Quality control. **a**, UMAP of 96h, 108h and 120h TLS coloured by predicted cell cycle, showing that cells do not cluster by cell cycle. To this end, first for each gene the cell cycle was estimated and subsequently this source of heterogeneity was regressed out so that the scaled data could be used for UMAP calculation without a cell cycle bias. **b,** Statistics for each of the cluster before correction for number of UMIs, genes and mitochondrial fraction. Cluster 10 is a clear outlier showing a strong bias towards an extremely high mitochondrial fraction and cluster 11 was marked by a high percentage of cells with an unusual number of UMIs and genes such that after removal of these cells, only a small fraction of cells was left. Both, cluster 10 and 11, were excluded from further analysis. **c,** Table displaying the average number of UMIs and genes detected for all cells separated by cluster. The average number of UMIs and genes is homogeneous between clusters and does not show any outliers. **d,** UMAP of *in vivo* cells from Somite and PSM cell states. UMAP coloured by initial cell state (left) compared to more stringent cluster assignment (right). The stringent cluster assignment results in four sub-clusters, that split PSM into two equally sized clusters, while the Somites split into one larger cluster and a second group of cells that are located at the transition of the previously identified Somites and PSM cluster. **e,** Heatmap of normalized expression of marker genes for the newly assigned four clusters coming from *in vivo* PSM and Somite cells. The two PSM cluster both show strong expression of the same subset of marker genes, however, the new small subset transitioning Somite cells has a clear anterior PSM (aPSM) signature. **f,** Correlation analysis of bulk RNA-seq, showing very high correlation between replicates of the same treatment. Only genes covered by all samples and expressed in at least one sample were considered (*n* = 29,963) and the pairwise Pearson correlation coefficient was calculated on the log_2_(TPM+1) values.

